# Standardised Measurements for Monitoring and Comparing Multiphoton Microscope Systems

**DOI:** 10.1101/2024.01.23.576417

**Authors:** Robert M. Lees, Isaac H. Bianco, Robert A.A. Campbell, Natalia Orlova, Darcy S. Peterka, Bruno Pichler, Spencer LaVere Smith, Dimitri Yatsenko, Che-Hang Yu, Adam M. Packer

**Affiliations:** Science and Technology Facilities Council, Octopus imaging facility, Research Complex at Harwell, Harwell Campus, Oxfordshire, UK; Department of Neuroscience, Physiology & Pharmacology, University College London, UK; Sainsbury Wellcome Centre, University College London, UK; Allen Institute, Seattle, USA; Mortimer B. Zuckerman Mind Brain Behavior Institute, Columbia University, New York, NY 10027, USA; Independent NeuroScience Services INSS Ltd, Lewes, East Sussex, UK; Department of Electrical and Computer Engineering, University of California Santa Barbara, USA; DataJoint Inc., Houston, TX; Department of Physiology, Anatomy, and Genetics, University of Oxford, Oxford, UK

## Abstract

The goal of this protocol is to enable better characterisation of multiphoton microscopy hardware across a large user base. The scope of this protocol is purposefully limited to focus on hardware, touching on software and data analysis routines only where relevant. The intended audiences are scientists using and building multiphoton microscopes in their laboratories. The goal is that any scientist, not only those with optical expertise, can test whether their multiphoton microscope is performing well and producing consistent data over the lifetime of their system.

## Introduction

Multiphoton excitation was originally described in 1932^1^ and the first scanning multiphoton microscope demonstrated in 1990^2^. Since then, the technology has been driven by the use of multiphoton microscopy for imaging deep into scattering tissues. Many research labs now use either custom-built multiphoton microscopes or instruments from commercial manufacturers. However, there are very few accessible tools and protocols for quantifying a multiphoton microscope’s performance. This quantification is essential for maintaining imaging performance over time, comparing instruments, and rigorously reporting experimental procedures to enable reproducibility.

We think quantification protocols are necessary for several reasons. First, as developers, early adopters, builders, users, and facility managers, we have realised that there is not always a consensus on which metrics are needed, nor agreement on best practices when performing the relevant measurements. We want to share practices that we think provide the most crucial measurements both efficiently and accurately. Such measurements can provide valuable diagnostic information about system performance, especially when characterisation is performed regularly. Second, we aim to further push the transition of multiphoton microscopy from a frontier technology to a routine tool, similar to confocal microscopy. Third, we hope this effort will contribute to more reliable comparisons of results within and across laboratories. Fourth, and finally, we aspire to engage manufacturers to specify their microscopes’ performance in similarly quantitative ways and develop better tools for such characterisation. Overall, we desire to push the field to improve data quality and rigour for the purposes of efficiency, open science, and reproducibility.

## Overview of the protocol

This document is a collection of related protocols, many of which can be performed independently, but some of which need to be performed before others. Guidance on testing frequency is provided in each section. We provide sub-protocols to characterise both the excitation and collection sub-systems of the microscope. For excitation, we cover how to carefully and simply quantify the laser power on sample, the excitation volume (i.e. point spread function; PSF), the field of view size and homogeneity, and a protocol for optimising excitation laser pulse width on sample. For collection, we provide a method for monitoring the sensitivity of detectors (often photomultiplier tubes; PMTs), as well as one for quantifying the photon transfer function of the microscope as a whole. All sub-protocols assume that the reader has all the components needed for a multiphoton microscope, and that they are already configured correctly and controlled by software. While more detailed characterizations can be made with specialised (and expensive) equipment, we designed the sub-protocols to require relatively simple, inexpensive and readily-available resources.

## Comparison to other protocols

Alternative characterisation efforts are underway; for example the QUAREP-LiMi initiative [https://quarep.org/] that aims to improve quality assessment and quality control for light microscopy. This effort, with widespread community support from scientists and manufacturers, has already produced protocols for general light microscopy^3,4^. Our goals are different in that our narrower focus is meant to provide a single, comprehensive characterisation of a multiphoton microscope.

There are many protocols on various aspects of quality control for confocal microscopes, many of which use similar scanning optics to those used in multiphoton microscopes (PSF only: Cole et al Nat Prot 2011^5^ & ^4^, quantification: Jonkman et al Nat Prot 2020^6^, reporting: ^7^). Our effort is similar to a recent protocol on “Strategic and practical guidelines for successful structured illumination microscopy” (https://www.nature.com/articles/nprot.2017.019^8^). However, these protocols do not cover key issues specific to the multiphoton case (but see recent publications regarding laser assessment^9,10^). For the nonlinear excitation that enables multiphoton microscopy, pulsed laser sources are required and thus additional factors such as laser pulse repetition rate, pulse width, and compensation for pulse broadening require consideration. Additionally, test samples must undergo efficient two-photon excitation without bleaching or burning, which makes some quantifications particularly challenging. Finally, it is important to characterise multiple aspects in a coherent fashion so our goal is to have a collection of related protocols.

## Limitations

These protocols are intended as a practical resource for the multiphoton imaging community. We are purposefully focusing on measurements that can be performed by users with turn-key systems, i.e. those without advanced optical metrology equipment. For example, we are not including a complete characterisation of the pulse shape and spectral properties, which would require an expensive optical device (an autocorrelator) as well as significant optical expertise to use it properly. Therefore, this protocol is not a complete characterisation of all aspects of multiphoton microscopy hardware relevant for designing such a system. In short, this is not a ‘build’ manual as there are already many such excellent resources like^11–15^.

We have executed these protocols on many different multiphoton systems and made the step-by-step instructions as generic as possible, however it is still possible that some measurements may be difficult to perform as described on some systems, depending on software control limitations.

## Laser Power at the Sample

### Introduction

Measuring and monitoring laser power at the sample is critical in multiphoton microscopy, and the laser power output is an easy-to-check indicator of instabilities in the microscope system. First, emitted fluorescence, often the signal of interest, is proportional to the average laser power squared, so small changes in laser power can result in large changes to your data. Second, exposing your sample to significant laser power can cause photo-bleaching and photo-damage, altering your sample, and measurements. Broadly, there are two types of laser-induced photo-damage that can occur in multiphoton microscopy^16^. The first is local heating of the area being imaged, which is linearly related to the average laser power. While this effect may be minor for normal imaging conditions, it should not be ignored, as it has been shown to alter the nature of biological samples^17–19^. The second type of photo-damage is photochemical degradation, such as bleaching or even ablation, which is nonlinearly related to the average laser power^16,20,21^. Therefore, knowing your laser power is essential for consistency between experiments, for minimising or eliminating photo-damage, and for monitoring the health of your imaging system.

### Laser power control

Before reaching the microscope scan head, the laser beam is routed through a user-controlled variable power modulator (e.g. a Pockels cell, motorised half-wave plate or acousto-optic modulator). The modulator may be a device that intercepts the laser on the airtable or it might be integrated into the laser enclosure itself. Attenuators typically use polarisation or diffraction to send a defined proportion of the beam into a “dump” to safely absorb excess light, whilst the remainder passes through the optical system and reaches the sample. When the user “changes laser power” they are often altering the ratio of power going to the dump versus the sample.

Uncalibrated control software will usually allow the user to set laser power along an arbitrary scale, such as 0% to 100%. What those values mean exactly will depend on the software itself, but most likely they simply map to the minimum and maximum of an analog voltage output that is fed to the controller electronics of the power modulator. It is common for modulators to respond non-linearly to the control input so laser power may well be a sigmoidal function of the “percent power” value. The maximum power may even occur well before “100% power” setting.

Some microscope control software allows the user to calibrate the on-screen laser control output such that it is mapped directly to laser power in watts (W), removing any nonlinearity introduced by the power modulation hardware. Other software has no facility for calibration and will always display power in arbitrary units. Here we describe the most generic process, which is to manually generate a table for converting percent power into a value in W.

### Microscope configuration

Microscopes can be configured and controlled in a variety of ways that can affect laser power measurements.

There are often substantial losses in laser power as the beam traverses the path towards the sample.

For example, it is not unusual for the objective alone to have transmission efficiencies of only ∼ 70%^22,23^. For this reason, the laser power metre is placed after the imaging objective to get the best estimate of power arriving at the sample. Note that laser power measurements will be dependent on the objective used, and are only valid for that objective. A different objective will likely have different physical properties, such as anti-reflective coatings, different number of glass elements, and a different back pupil diameter, all of which will affect transmission and therefore laser power at the sample.

There are two ways of measuring the time-averaged laser power under the objective: either with the beam stationary (typically centred in the imaging field of view) or with the beam continuously scanning. Where possible it is preferred that measurements are made with a stationary beam or scanning with beam blanking (see **Fig. 1A** for explanation) disabled, as this measurement is independent of changes in effective duty cycle of the scanning, and easily comparable across different instrument and power meters. On the other hand, during normal scanning the beam blanking is enabled, reducing the duty cycle and leading to a lower average power measurement. Measuring in this way more accurately represents the average power deposited on the sample under those exact scan conditions (see **Fig. 1A**). If the aim is to strictly monitor laser power for internal comparisons only, then it is sufficient to simply use one method consistently. However, a stationary beam is preferred if you want to compare externally (different microscopes) or report your laser power in publications, as all other calculations of power density can be made from the stationary measurement e.g. mW/mm^2^ if the scan parameters are known.

**Figure 1.**
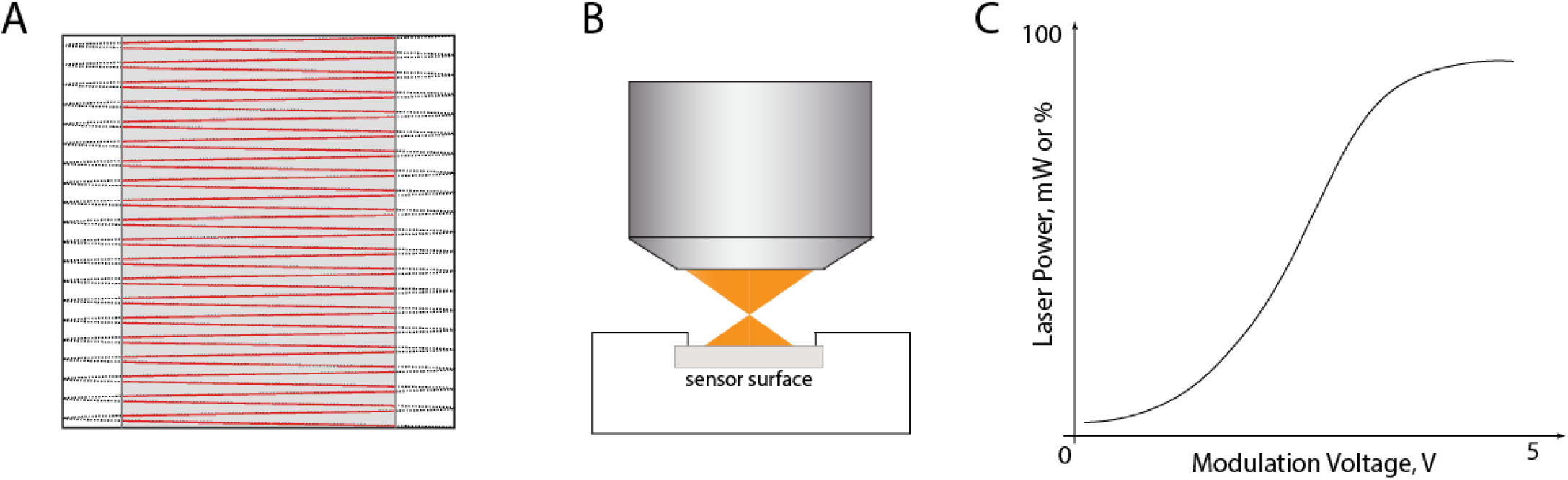
Laser power measurement. **A**. The path of a scanned focused laser beam over a sample. The beam moves sinusoidally along the fast axis (X in this case) whilst being scanned in orthogonal Y direction with a linear galvo. The area over which the beam moves is known as the scan field. On the left and right edges the beam slows as the scanner changes direction and turns around. In these areas the potential for photodamage is greatest, as the beam is travelling more slowly over the sample. Thus the beam is typically “blanked” or disabled during these epochs. In a resonant scanning microscope the beam is usually blanked about 30% of the time. The image field (red lines and grey region) is the area over which the beam is on and capable of exciting fluorescence. The dotted lines indicate the blanked turn-around regions. A power meter cannot distinguish these states and so returns an time-averaged power value over the whole scan field if the microscope is scanning during a measurement. **B**. Position of the power meter head with respect to the laser beam exiting the objective. The sensor surface should be close but must not be at the working distance of the objective lens, as the focussed beam may damage the sensor surface leading to unreliable measurements and permanent damage to the sensor. **C**. Example laser power calibration curve. Output power can be represented as a percentage of total available laser power, or in direct power units, mW, depending on microscope configuration. The purpose is to create a lookup table that allows linear adjustment of power on the edges of the modulation range for modulation devices (like Pockels cells) that have non linear response.

### Time-averaged laser power measurement

This protocol describes the measurement of time-averaged laser power at the sample (after the objective lens). However, the average power is not the complete picture. Lasers used in multiphoton microscopy are pulsed, which can increase the efficiency of multiphoton excitation^24^. The power during a pulse is known as the peak power, which is often orders of magnitude higher than the average power. The peak power can be easily estimated from the average power under the simplifying approximation that the pulse envelope (the “shape” of the intensity profile of the laser pulse) is rectangular:

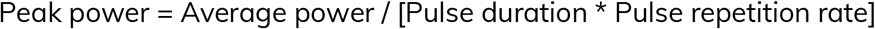

You should consider the peak power as well as average powers when comparing experiments on different microscopes and lasers. Two systems with the same average power could have vastly different peak powers, leading to different amounts of photo-bleaching, emitted fluorescence and photo-damage.

The aim of this protocol is to equip anybody with a simple metric for monitoring the laser power reaching the sample on their multiphoton microscope, the time-averaged power. This protocol can not be used to measure actual peak power, since this requires knowledge of the pulse duration (see Grouped Delay Dispersion Optimization), and pulse envelope, which is a much more complicated measure.

Time-averaged laser power is measured with either a photodiode or thermal power sensor. Photodiodes convert light directly into an electrical signal, whereas thermal power sensors convert the thermal energy deposited by the light into a measurable voltage. Thermal power sensors are often recommended because photodiode sensors can saturate depending on the peak powers of your laser, though thermal sensors are slower to respond to power changes, and more sensitive to external conditions. The wavelength range, power range, sensor area and sensor resolution should all be taken into account when choosing a power sensor. The wavelength and power range should match the lab’s imaging needs. The sensor area should be larger than the exit area of your objective to allow sufficient light to get to the active sensor area; a large objective lens may require a larger opening (e.g. Thorlabs S425C). Finally, the sensor resolution must be fine enough to give you the accuracy you need (e.g. 1 mW or 0.1 mW depending on applications).

### Materials

#### Equipment

- Laser power meter
- Compatible power sensor head
- Suitable clamping hardware or adhesive (tape, sticky tack) to ensure the sensor remains stationary during measurements
- Objective lens(es) used for imaging

### Equipment setup

1. Turn on the microscope hardware necessary for controlling laser beam scanning and laser shutters.
2. Turn on the laser and select the wavelength normally used for imaging applications. (WARNING: Lasers can emit light that can be hazardous to people and property - it is critical to have have adequate provisions in place to prevent unwanted laser exposure - local laser safety officers can provide guidance) OPTIONAL: Some laser modulators, such as Pockels cells, are temperature sensitive. If using such a modulator, open the required shutters so that the beam passes through the device to warm it to an equilibrium temperature. In an ideal setup, there will be a hard shutter downstream of the Pockels cell, and this shutter is used to prevent light from entering the microscope when the beam is “off”.
3. Allow the laser power output to stabilise (at least half an hour, but preferably an hour). NOTE: You can do repeated power measurements during laser warm up to determine how long your system takes to stabilise. OPTIONAL: If using a Pockels cell, ensure the bias voltage is set to the value when it was last aligned.
4. Ensure the objective is clean. If not, clean the objective lens using a suitable method depending on the type of objective, and the contamination. Consult manufacturers recommendations if unsure of the procedure.
5. Install the objective in the microscope.
6. Plug the sensor connector firmly into the power meter console.
7. Turn on the power meter, and ensure the battery is charged or it is plugged into a power source to last for the duration of the measurements.
8. Set the power meter console wavelength setting to match the laser wavelength selected in step 2.
9. Place the sensor under the objective and secure the sensor in place using a clamp, bolt or other suitable adhesive for your setup. OPTIONAL: Use objective immersion media if the power sensor is designed to allow for it.
10. Centre the objective over the active area of the sensor in XY and set the height such that the laser will not be focused on the power meter’s sensitive surface. NOTE: It is a balancing act to set the objective-sensor distance. The sensor should be close to, but not at, the objective’s working distance, as this will ensure the sensor captures all light whilst also minimising the chance of damaging it with a tightly focused beam (**Fig. 1B**). More consistent readings will be obtained if the beam fills a large proportion of the sensor. Always take readings at the same distance from the objective to avoid variation in results.
11. ”Point” or “centre” the beam in the software, so it is stationary in the centre of the field of view. If this is not possible, a similar result is achieved by zooming in by a factor of 10x or 20x and scanning a small area with beam blanking disabled.

### Procedure

CAUTION: Ensure that local laser safety rules are followed at all times to keep you safe from fire and damage to eyes/skin.

The way power readings will be taken will depend on your hardware and software. There are three possibilities:

- Manual readings: the user must set the power level in software and record the laser power value reported by the power meter console.
- Software calibration: a calibration process provided by the microscope acquisition software. Note: the calibrated power in microscope software may not be accurate for low values. If you need low values, such as 5 mW, then you should check the correct percent power manually. Calibration curves are wavelength-specific because laser max power and Pockels cell bias voltage both vary with wavelength.
- Custom software: a custom software tool that reads values from the power meter console whilst controlling laser power, i.e. a Python script.

1. ”Zero” the power meter.
2. Ensure the laser is set to the desired wavelength.
3. Open any final shutters so that the beam will pass through the system to the sensor.
4. Find the minimum power by adjusting the software’s power modulation control until a minimum reading is seen on the power meter.
5. Record the power meter measurement (usually in units of mW) and power modulation value (i.e. %, volts, arbitrary units) for this minimum power.
6. Find the maximum power by adjusting power modulation until a maximum reading is reached. NOTE: Do not exceed the power sensor’s upper limit, as this may damage the sensor or lead to inaccurate readings.
7. Record the power meter measurement and modulation value for this maximum power.
8. Record intermediate laser powers and modulation values that are commonly used in your experiments.
9. Arrange the above power meter measurements and modulation values in a table for future comparisons. Provide information regarding how the reading was made (stationary or scanning beam) and if scanning, report the temporal fill fraction. If you have access to different power meters, report the model number or serial number of the sensor head.
10. If using more than one wavelength, repeat steps 2-9 for those.
11. Refer to historical power measurements and ensure that the laser power is not changing. If it is, try to determine the source of the variation by following some troubleshooting steps below.

### Troubleshooting

A change in laser power can be the result of many different factors. Below, we suggest some things to consider when troubleshooting. If you are not trained to carry out alignment or adjust table optics, contact an experienced individual in the lab or department, or contact a microscope or laser engineer.

#### Laser throughput is degrading

Lasers will not always output the same power; over time they may degrade. Some lasers include internal monitors that allow you to measure the power inside the laser head. By logging these periodically you can determine if the laser power output is reduced.

#### Laser alignment has changed

Laser power at the objective is highly sensitive to beam alignment. For instance, if the beam is not hitting the scanners correctly, it can be partially clipped or blocked. Measurements should be repeated in case laser alignment is adjusted.

Record the laser power at different points in the beam path to determine where power is being lost. Making logs of these values at the different points along the path after each alignment change will make troubleshooting later easier as you can compare to these values and find where misalignment may be. Useful locations to log transmission would be before and after the modulation device as well as before and after the scanhead.

Having alignment targets in two or more places on the table and one at the objective mount is also a simple way to troubleshoot if measurements are off due to alignment. If the beam is not hitting the centre of a target, it suggests the laser is misaligned before that point. Adjustable aperture irises are convenient for this purpose, as they can be mounted directly in the path, yet opened for free beam passage under normal conditions.

#### Laser power value changes over the course of minutes and is not stable

You might notice minor (< 5%) laser power drifts during the day. “Fast” or larger drifts should not occur however - for example a change of 5 or 10% over the course of 15 minutes. If the laser itself is stable, such a change could be due to instability of the power modulator. Pockels cells in particular are temperature sensitive, and a unit with an internal or directly affixed beam dump can drift significantly as the dumped beam could cause strong local heating, especially for lasers with power outputs of a few watts. In this case, the beam dump should be decoupled from the Pockels cell, with a short distance between them. [CAUTION: beam dump repositioning should only be attempted by a competent, trained, individual. The discarded beam exiting a Pockels cell is of high power and often exits at a dangerous angle, making an eye-strike possible.] Similar temperature instabilities can occur if the Pockels cell is downstream of a shutter and only gets the laser beam when imaging is active, as its internal temperature will change as the laser passes through the cell.

Another potential source of apparent power variation is improper use of thermal sensor heads. Thermal power sensors are sensitive to ambient temperature and physical handling, especially at lower power values. The sensor should be allowed to equilibrate and only used once zero drift has ceased.

### Power sensor has not been calibrated recently

Power sensors may need calibration every 2-3 years, if your values are slowly changing, you may want to recalibrate your power sensor.

## Field of View Size

### Introduction

In conventional laser scanning systems, the image is created by scanning the focused laser beam over a rectangular area with a set of mechanical scanners that steer the beam in X (fast) and Y (slow) directions. This scan pattern is known as raster scanning (**Fig. 1A**). The beam travels over the rectangular area in the tissue, exciting fluorophore molecules along the path of the scan. The resulting fast fluorescence emission (often a few nanoseconds) is detected and integrated over a short duration (the pixel dwell time) to create an intensity value for a single pixel. Consecutive pixels are captured and used to create an image. The area on the sample covered by the scan is known as the field-of-view (FoV). The maximum area possibly scanned is limited by the optics and hardware, but under normal conditions is restricted by controlling the scanners’ scan angle. This can be set in software by altering the scanner’s amplitude to “zoom in” on some specific structure, or see the maximum FoV.

Knowing the physical size of the FoV allows the calculation of the pixel-pitch in microns. From this you can measure physical dimensions of features in the imaged sample as well as select the correct zoom factor to achieve spatial sampling that takes advantage of the full optical resolution of the system (Nyquist sampling), if required. Furthermore, measuring FoV size is an essential step for characterization of the imaging system: measurements in other sections of this protocol (e.g. Spatial Resolution, FoV Homogeneity) will rely on an accurately calibrated FoV. An uncalibrated system may have non-square pixels; knowing the pixel size is a prerequisite for correcting this should it be necessary.

The maximum FoV of an imaging system is defined by the maximum scan angles in the X and Y dimensions and the excitation optics, but the working field of view used in experiments should be selected based on other characteristics, such as field homogeneity (see next section), field curvature, and distortion (see Troubleshooting section). The FoV size is also used in mosaic imaging where tiles of images are acquired in a grid and merged together. FoV size is not expected to change from year-to-year, so monitoring its size is not required unless the optics or scanners in the excitation path of the microscope have been changed. Here, we describe how to measure the maximum effective FoV of a laser scanning multiphoton microscope.

We will measure physical dimensions of the imaged FoV at minimum zoom factor (the largest possible scan angle) and evaluate field distortion based on that image. For this, we will acquire an image of a fluorescent sample with features of known geometrical dimensions and convert this to microns in the X and Y axes. The sample can be a microscopy grid overlaid over a Chroma fluorescent slide, or a copper grid used in electron microscopy. Copper grids are cheap, will emit light when stimulated with a multi-photon laser and are robust to damage. They can also be overlaid on top of a Chroma slide to create a negative image.

### Materials

#### Equipment

- Calibration sample - one of the following two options can be used:
  1. Grid slide (Thorlabs part number R1L3S1P or Edmund Optics part number 57-877) overlaid over an autofluorescent plastic slide (Chroma part number 92001). To prepare such sample, follow these steps:
    a. Clean all dust and dirt from both slides.
    b. Place the autofluorescent plastic slide horizontally.
    c. Identify which side of the grid slide has etchings.
    d. Add a droplet of water onto the chroma slide and place the grid slide such that the etchings face the autofluorescent slide.
    e. Clamp two slides together with a mechanical clamp, adhesive or epoxy resin.
  2. Copper grid used for electron microscopy (e.g. SPI supplies, part number 2145C-XA). These grids have a pitch of 25 microns with 19 micron holes. Dry copper EM grids will emit light when stimulated by a multiphoton laser. Prepare such a sample with the following steps:
    a. While keeping the grid dry, place it on a conventional glass slide or an autofluorescent slide (if a bright negative image of the grid is needed).
    b. Place a coverslip over the top of the grid.
    c. Seal the coverslip with nail varnish.

### Equipment setup

1. Turn on the laser.
2. Set the laser wavelength to one used for imaging experiments.
3. Start the microscope control/image acquisition software.
4. Install a microscope objective lens used for imaging experiments.
5. Place the sample under the objective lens as perpendicular to the optical axis as possible to eliminate any tilt with respect to the imaging plane. NOTE: it is possible to use the microscope’s laser beam to align the sample to the objective. For this, lower laser power such that no more than a few mW exits the objective. Close any iris in the excitation path to only let through the very centre of the beam. Make sure the beam hits the sample and then use an IR viewing card and an IR viewer to visualise the reflected beam. Here, the reflection will be very weak, on the order of 1-5%, so an IR viewer is needed to visualise it. Adjust sample position such that the reflected beam propagates directly along the incident beam.
6. Apply any immersion media appropriate for the objective lens and position the lens at roughly its focal distance away from the sample.

### Procedure

1. Focus on the calibration sample gridded surface. If using a fluorescent slide overlaid with a grid, image the very surface of the fluorescent slide to get a negative image just underneath the grid (**Fig. 2B**).
2. Acquire an image of the calibration sample at the minimum zoom to give the largest FoV (**Fig. 2A, 3A**). NOTE: For large FoV microscopes (>1 mm) it might be difficult to eliminate tilt completely, in this case it’s necessary to acquire a z-stack that covers the axial extent of the tilt. Similarly, a z-stack might be needed when the system has field curvature.
3. Process the acquired image to determine the FoV and pixel size.
  a. Count the number of grid squares along a row or column of the grid across as much of the FoV as possible.
  b. Make a note of that distance in pixels for calculating the micron-to-pixel conversion.
  c. Multiply the number of grids by the grid pitch size in microns to give the total length in microns.
  d. Divide the distance in pixels by the distance in microns to get the micron-to-pixel conversion and multiply that by the total pixels in the image to get the FoV size.
  NOTE: As counting the lines of a copper EM grid is time consuming, using an automated tool may be preferable (see https://github.com/SWC-Advanced-Microscopy/measurePSF, tool: Grid2MicsPerPixel, **Fig. 2C**).
4. If the microscope has other imaging paths, repeat the measurement using these imaging modalities (e.g. epifluorescence or brightfield, **Fig. 3A**).
5. Evaluate whether there is pincushion or barrel distortion^25^ in the images (**Fig. 2B**). Presence of these aberrations would signify an off-axis optical performance degradation in the system.
6. Comparing images acquired in multiphoton and widefield modes allows to characterise rotation/mirror effects between the two imaging modes (**Fig. 3**).
7. Repeat steps 1-6 for any other objectives used for imaging, as calibration is only relevant for the objective used to do that calibration. Calibration files are usually saved for each objective in the acquisition software so that the pixel size is calibrated for every acquired image without any manual metadata editing.

**Figure 2.**
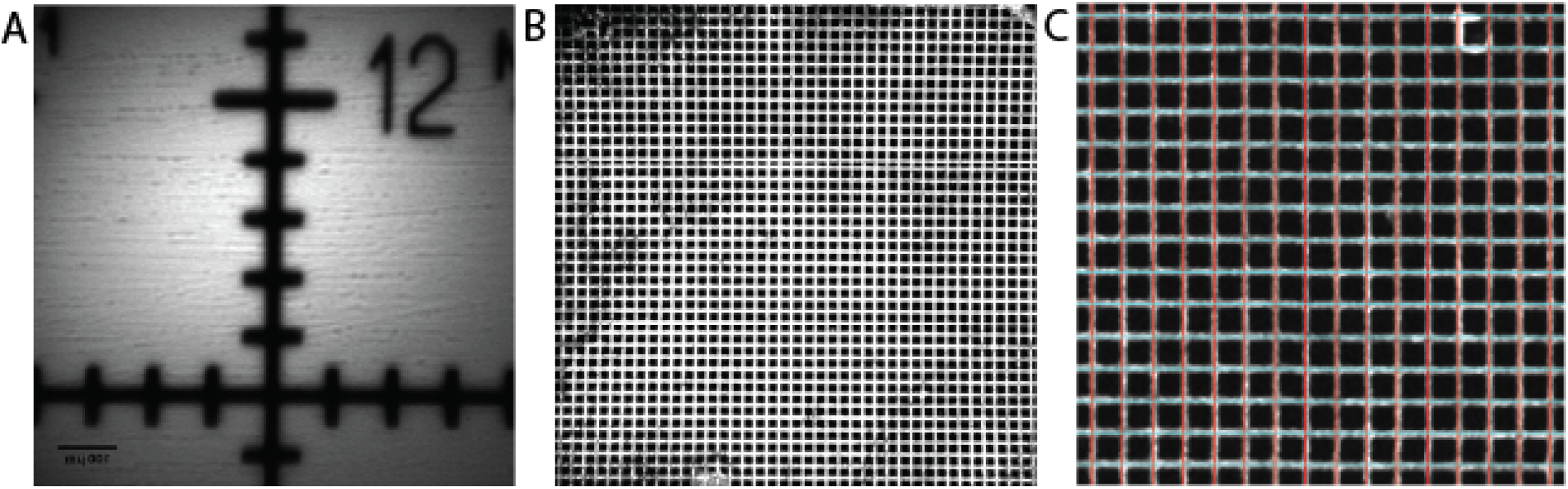
FoV size measurement. **A**. Two-photon image of a 1mm grid with 100 um divisions for maximum scan angle (minimum zoom). Here, the FoV size is ∼ 800 um. **B**. Image of a 25 micron copper EM grid. This image shows the ∼ 1200 um FoV and displays pincushion distortion at the left and right edges. **C**. Grid lines are detected and overlaid on top of the EM grid image using the “Grid2MicsPerPixel” software tool.

**Figure 3.**
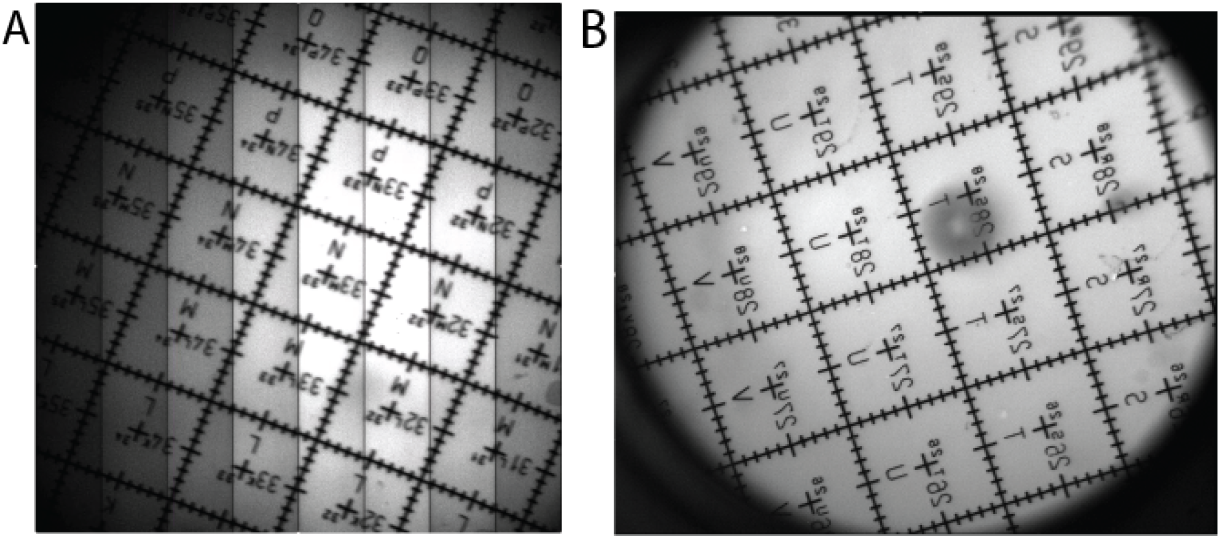
FoV Size comparison for two-photon and epifluorescent modes on a large FoV microscope (Mesoscope). A. Tiled two-photon image of the Mesoscope FoV (∼5000 um). **B**. Epifluorescent image of the Mesoscope FoV showing a ∼ 45 degree rotation, and vertical reflection, compared to the two-photon image.

### Troubleshooting

When assessing images acquired in this part of the protocol, pincushion/barrel distortion and z-plane (field) curvature might become apparent. Pincushion and barrel distortion are usually caused by optical aberration at high scan angles in excitation optics. Ideally, the XY beam displacement in the imaging plane depends linearly on the scan angle. Practically, due to optical design compromises, this linearity breaks down at higher scan angles, usually towards the corners of the image. Unfortunately, this is a limitation of the optics and is most easily resolved *post-hoc* by correcting the images. For applications such as functional *in vivo* imaging it may not be necessary to correct for distortion.

For systems that generate larger images by moving stages and for resonant-galvo-galvo (RGG) systems capable of tiling single FoVs into a mosaic of multiple scanned areas, one needs to calibrate the XY stage/scanner step to achieve proper mosaic FoV. This is usually performed after measuring the FoV size. Software like ScanImage provides an interface where the operator can iterate a few times to achieve correct tiling.

## Field of View Homogeneity

### Introduction

Field of view (FoV) homogeneity is a metric that defines the system’s uniformity of excitation and collection of light across the FoV of the sample. Measuring field homogeneity is easy to do and can be used to understand the data quality across the image, and also to know the experimental FoV size that minimises this variation. Large structured variations in homogeneity can also highlight potential problems with microscope alignment or debris on optics in the path.

One common inhomogeneity that can be revealed from this measurement is known as “vignetting”, where signal is reduced at the edges of the FoV. Vignetting is important to consider when choosing the working FoV for experiments as reduced signal at the edges of the FoV will cause a decrease in signal-to-noise.

Vignetting is unavoidable when using large scan angles that are at the design limit of the microscope hardware and optics. The limits are defined by the optical properties of the microscope, in particular the objective lens. Manufacturers usually specify the FoV size over which an objective’s performance is said to be “diffraction limited”. Beyond the specified FoV size, the objective’s ability to focus light degrades and causes vignetting and other optical aberrations. Typically the objective is the principle limit for the FoV size, but an improperly designed excitation or collection path might become a limiting factor depending on PMT sensor size, angle of acceptance, and the collecting lenses (for more on this topic, see ^26,27^).

This protocol describes how to assess the homogeneity of the field of view by imaging a uniform fluorescent sample and measuring how the image intensity varies across the image. The resulting measurements are a description of the overall system’s performance, meaning that any drop-offs in intensity could be caused by optical degradation in either (or both) the excitation and collection paths.

Many objectives used for *in vivo* multiphoton imaging have field curvature, where the imaging plane is not actually a plane, but is instead shaped like a shallow bowl. This is particularly noticeable for some large FoV systems^28–30^. To eliminate the effect of small field curvature on field homogeneity, the sample for this measurement should be sufficiently thick that the beam remains in the sample across the entire field. Either a fluorescent bath (e.g. fluorescein dissolved in water 1:10,000) or a thick autofluorescent slide (Chroma part number 92001) are good choices.

## Materials

### Equipment

- A uniform fluorescent target (either a fluorescein bath or thick fluorescent plastic slide)

### Equipment setup

1. Turn on the laser and allow it to stabilise (usually between 30 minutes and 1 hour).
2. Set the laser wavelength to one used for imaging experiments.
3. Start the microscope control/image acquisition software.
4. Install the microscope objective lens to be used for imaging experiments.
5. Place the sample under the objective at the objective’s working distance (usually written on the lens, or found in manufacturers specifications).
  a. If the sample is a fluorescein bath:
    i. Prepare the bath by diluting fluorescein 1:10,000 in water inside a suitable petri dish.
    ii. Immerse the objective tip in the bath.
  b. If the sample is a fluorescent plastic slide:
    i. Place the slide under the objective at its working distance and use an appropriate immersion medium.

### Procedure

#### Homogeneity measurement

1. Set the laser power down to 5 mW (for an 80 MHz pulsed laser, for considerations see Laser Power on Sample).
2. Begin imaging and slowly turn the PMT gain up from 0 until signal from the sample is seen to avoid damaging the PMTs with a potentially very bright sample. If no signal is seen, adjust the objective focus up and down to search for signal from the sample before turning up the PMT gain further.
3. OPTIONAL: If using a fluorescent slide, find the sample surface then slowly lower the objective until all hints of the surface inhomogeneities have vanished. You will probably need to go down about 100 to 200 microns.
4. Acquire an image of the uniform sample using the largest FoV size the system allows for. The number of pixels needs to be sufficient to capture the majority of the inhomogeneity. Additionally, an average of multiple images is advisable to reduce noise.
5. If this is the first time you are performing the measurements, acquire more images using higher zooms (smaller FoVs). For example, 50% and 25% of the full FoV. This will help diagnose inhomogeneities that could arise from the laser power modulator (see Troubleshooting).

#### Analysis

5. Use FoV size and aspect ratio data acquired prior (see Field of View Size) to translate pixels to microns.
6. Normalise the image by dividing all pixels in the image by the maximum pixel value.
7. Plot a line profile through the horizontal/vertical axes and through the diagonals of the FoV (**Fig. 4**). The horizontal/vertical profile is informative about each scan axis (from the X or Y scanner).
8. Evaluate how symmetrical the brightness across the FoV is. Because of the radial symmetry of most optical systems, a well aligned microscope should have the brightest part of the field in the centre of the image and the brightness should fall off evenly around the centre.
9. If previous measurements have been made, refer to them to determine if there have been any changes in homogeneity that need addressing.

**Figure 4.**
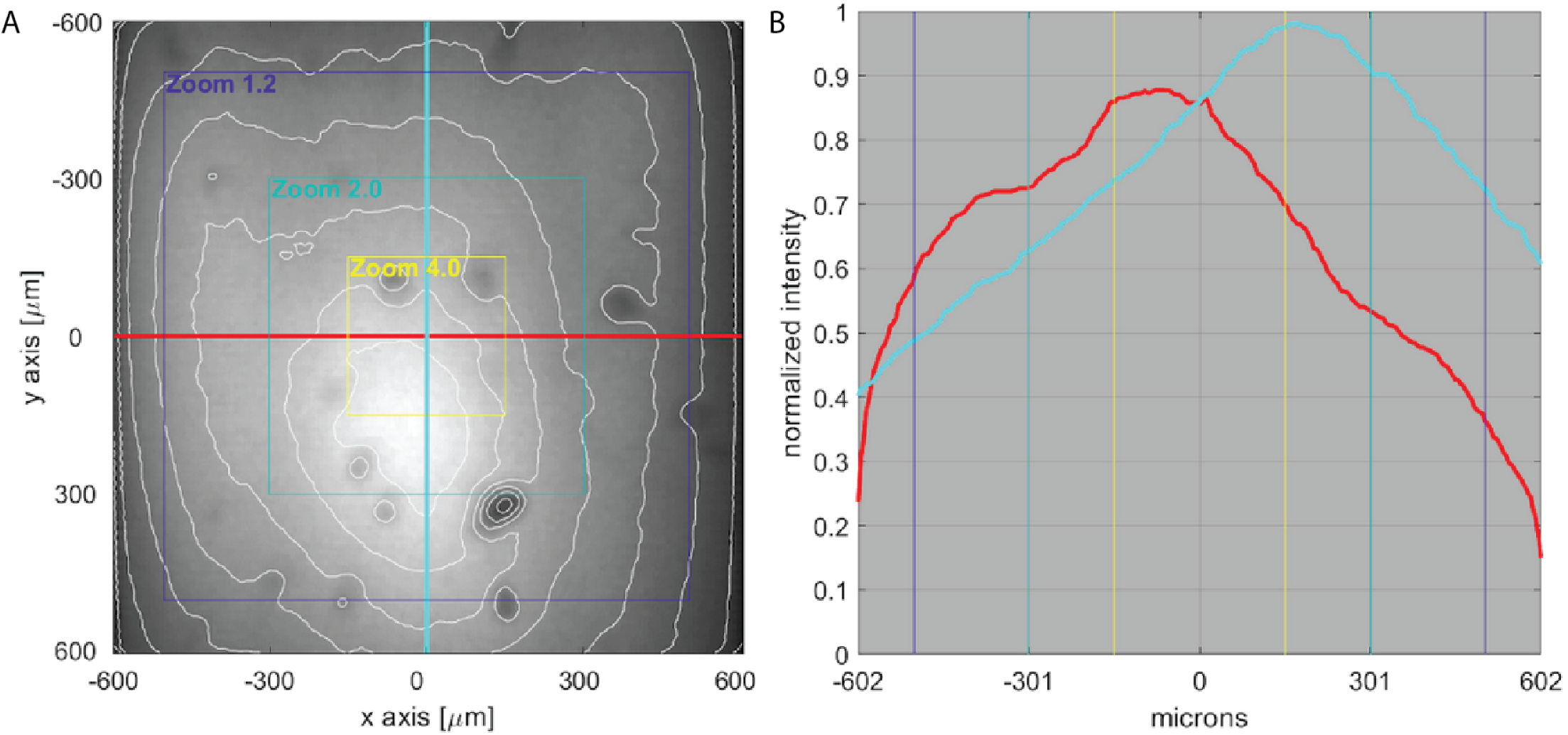
Field of view homogeneity. **A**. Homogeneity calibration image of the uniform fluorescent slide acquired using NIkon 16x objective at maximum scan angle on a system that allows for large scan angles. Homogeneity drop-off profiles and different zoom factor overlays are shown (different systems may have different scan areas for the corresponding “zoom” factor). The area of peak brightness is offset downwards slightly along the rows. This indicates a possible misalignment in the optics. Dark spots arise from imperfections in the slide surface. **B**. Intensity profiles along horizontal (red) and vertical (cyan) axis demonstrate non uniformity of excitation at the maximum zoom factor.

### Troubleshooting

The short focal length of the microscope objective means that alignment at this point is critical. To achieve guaranteed optical performance of the objective lens, the incoming laser beam must be collimated and centred on the objective. To minimise vignetting, the beam must optimally fill the objective’s back pupil, which must be properly conjugated with the scan mirrors. If the beam is not stationary at the back pupil during scanning, vignetting will be more pronounced.

#### Pupil conjugation issues

There are several observations that can come up during this measurement that are useful to troubleshoot the health of the system. Firstly and most commonly, if the intensity drops off very fast towards the edges, it usually signifies problems of the pupil conjugation in the system - to check for that one needs to evaluate whether the beam “walks” at the back pupil.

#### Laser alignment is not optimal

Asymmetric drop-off of the fluorescence towards the edges, or off-centre peak usually point to the problems with alignment, this can be confirmed with the measurement of spatial resolution (see Spatial Resolution). The geometry of the excitation volume will most likely be less tight in the area of the FOV that appears darker.

#### Problems with the sample

Local inhomogeneities of the image usually point to either bubbles in the immersion media, or inconsistent fluorescence in the imaging sample. If using a slide, try cleaning it.

#### Presence of dirt on optics

There may be some dirt on optics somewhere in the path, it is not unheard of for dust to burn on to optics due to high laser powers. Try using compressed air to dust the optics in the path to identify if any are causing issues, or where possible directly observe lenses to check for burnt on dust.

#### Pockels cell ringing

One potential source of image inhomogeneity that is not related to the optical properties of the microscope is intensity “ringing” that originates in a Pockels cell (**Fig. 5**). Some microscopes use an electro-optical power modulator known as a Pockels cell to rapidly control laser power through the application of large voltages to the Pockels cell. The materials used in these cells can have a strong piezo-mechanical effect, resulting in a damped oscillation at the cell’s resonance frequency. This is typically seen on the fast scan axis, on packers cells that do not have any additional damping (the so-called BK “clamped” option on one common brand, Conoptics). The resonance manifests as an unusually shaped inhomogeneity that extends along the microscope’s fast axis and does not change shape with zoom. Disabling beam blanking should reduce or eliminate the problem if it originates from the Pockels cell. It is for this reason that we recommend *initial* vignetting measurements are done over a range of zooms. Note that Pockels Cell resonance impacts on inhomogeneity may look different to that shown below, as they will depend upon the specifics of how the sample is imaged, and the nominal scan line duration.

**Figure 5.**
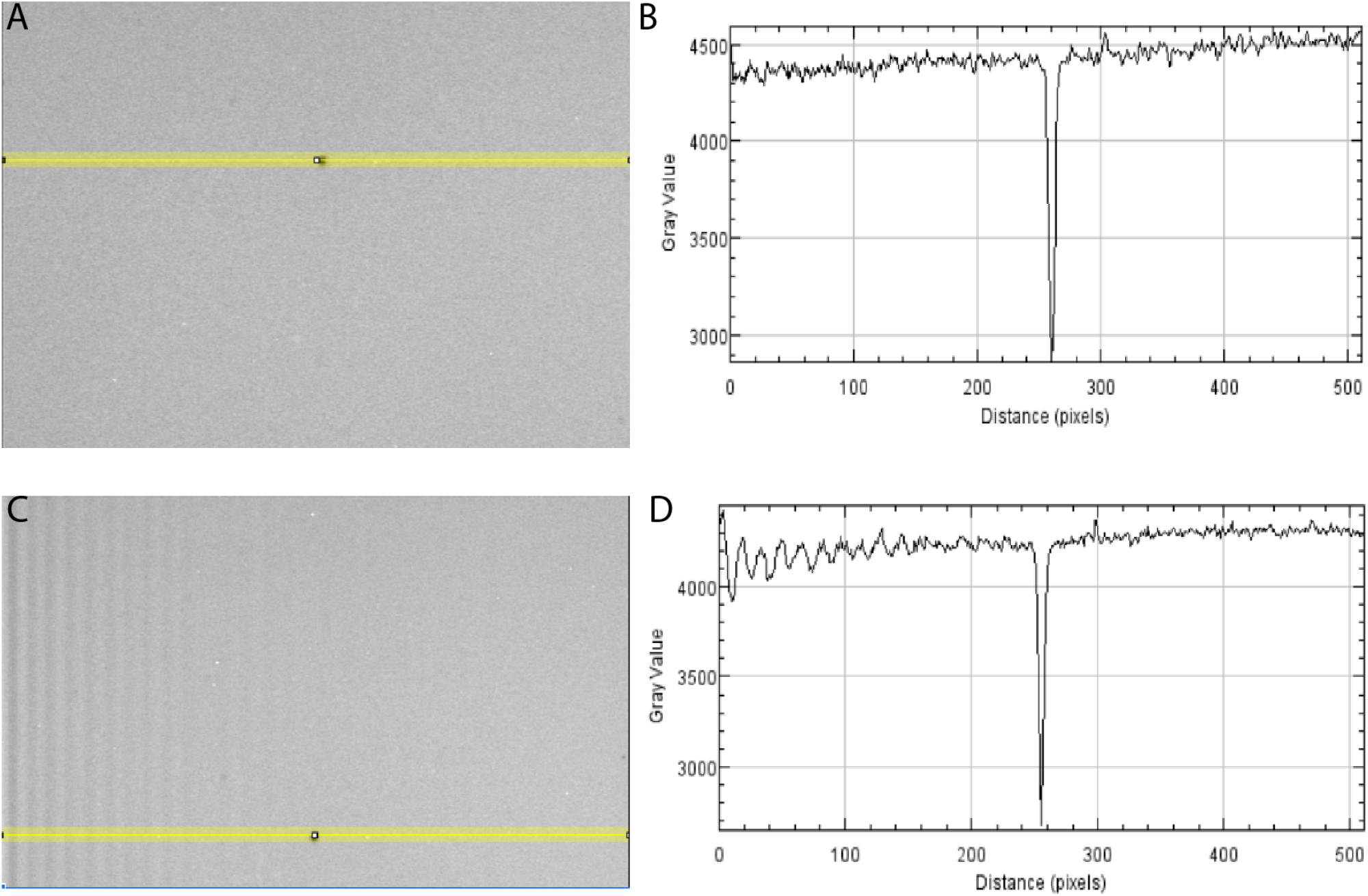
Pockels Cell resonance effect. **A**. Image of a homogeneous fluorescent medium with no ringing effect. **B**. Line profile along the yellow line in (A) that only shows a drop at the dark spot along the line. **C**. Same image as A, but with Pockels cell ringing visible on the left side. **D**. Line profile along yellow line in C that shows intensity oscillations on the left side of the image where Pockels cell ringing is present. This effect does not change with zoom factor. The line profile is chosen to extend over a darker spot to highlight the magnitude of the ringing.

### Spatial Resolution

#### Introduction

##### Excitation volume

Laser scanning imaging systems such as multiphoton microscopes provide spatial resolution governed by their point spread function (PSF). The point spread function is the impulse response of an imaging system. In other words, the PSF describes how a point object (an “impulse”) in the object space will be “spread” by the imaging system, and thus appear in the acquired image. The PSF is a three-dimensional spatial function whose shape can vary depending on the focusing optics of a system, and across the field-of-view. Moreover, nonlinear excitation and excitation saturation can influence the effective spatial resolution in multiphoton microscopy ^31^. Thus, in this protocol we will refer to the excitation volume geometry to specifically refer to the shape of the volume that is excited by the focused laser light at a given point in time. The geometry of the excitation volume governs how the smallest features will appear in the microscope and the effective spatial resolving power of the microscope. What is commonly characterised in multiphoton imaging is the shape of the excitation volume, often approximated by the FWHM of the axial and radial profiles, and that is the process we describe in this protocol.

##### Diffraction-limited or not?

The measurement of an excitation volume matters because it indicates how an imaging system resolves signals from structures in the sample, and the profile of the excitation volume is often engineered depending on the requirement of the application. For common use cases, the imaging system is designed to achieve a diffraction-limited PSF (the highest resolution possible for a system limited due to the physics of diffraction), which would ideally be smaller than the structures of interest being observed ^31–33^. In this way, the emission from the excitation volume is dominated by signals from individual structures, and contributions from neighbouring structures are minimal. We note that the term “diffraction-limited” is typically applied to beams with planar or gaussian wavefronts that then enter and are focused by standard objectives) In contrast, there are purposefully designed non-diffraction-limited PSFs, which are matching or much larger than the structure of interest, are utilised to rapidly sample volumes of tissue, especially when staining is bright and sparse^34–39^. Regardless of the experimental approach, excitation volume geometry should be characterised and monitored over time to ensure consistent resolution within a set of experiments. Although the example shown in this protocol is the diffraction-limited case, this protocol is empirical and applies in non-diffraction-limited cases as well.

##### Factors affecting the excitation volume

The numerical aperture (NA) is a unitless parameter that characterises the range of ray angles that an optical system can accept or emit. The NA and wavelength of the excitation light are the major factors that influence the smallest possible spatial extent of the PSF, such that larger NAs and shorter wavelengths lead to smaller PSFs (**Fig. 6**). The maximal NA is set by the objective, however, if the excitation light entering the objective underfills the back pupil, then the effective excitation NA will be reduced and the actual PSF is larger than the theoretical, diffraction-limited prediction for that objective ^40,41^. The profile of the beam used to illuminate the back aperture is approximately Gaussian in most cases, and its width is commonly characterised by the spatial extent over which the intensity exceeds 1/*e*^2^ of its maximum (*e* is the base of the natural logarithm). Thus, to use the full NA of the objective requires overfilling of the back pupil of the objective leading to some power being discarded.

**Figure 6.**
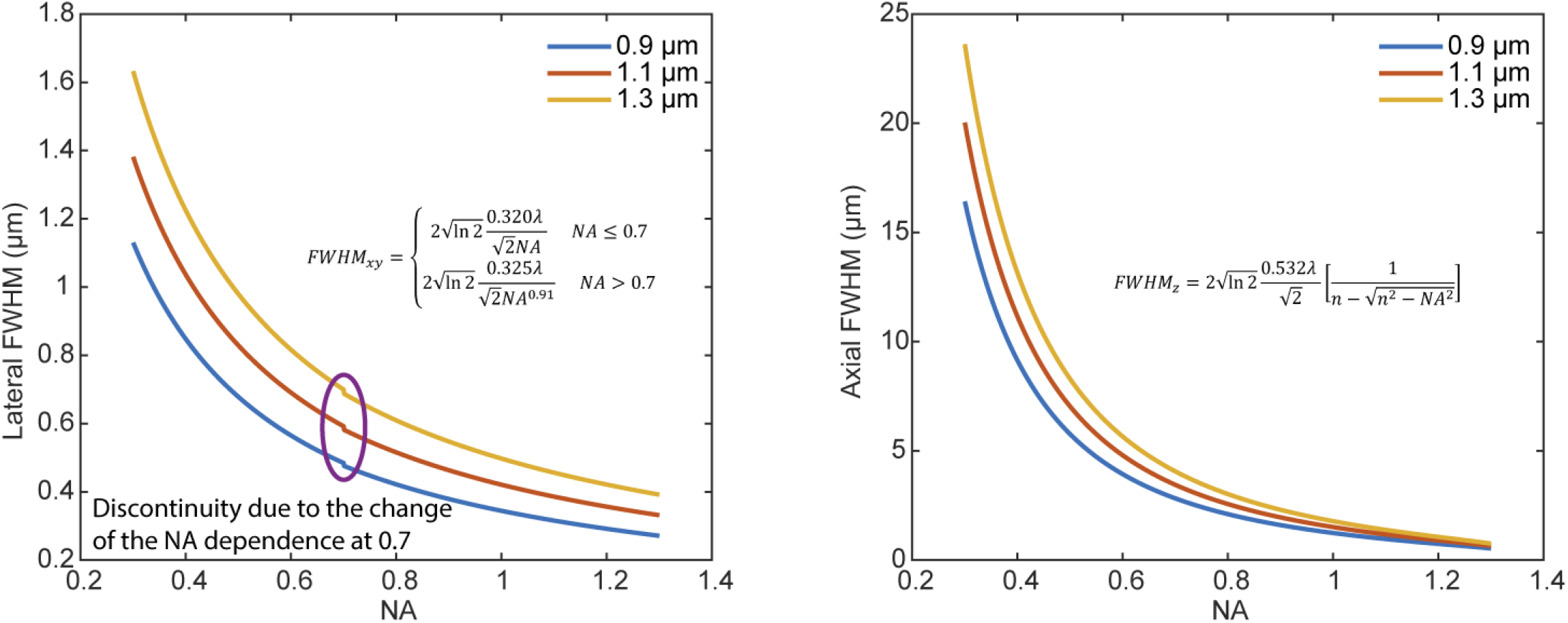
Two-photon FWHM as a function of NA and wavelengths^31^. The formula is adapted from Zipfel et al. 2003^31^. NA is the numerical aperture of the objective lens. In FWHMz, n is the refractive index of the medium where the sample is embedded, and set as water in this plot. The refractive index of the water, n, is wavelength dependent, and is [1.328, 1.3255, 1.3225] at [0.9, 1.1, 1.3] μm, respectively.

Sometimes objectives are deliberately underfilled to transmit more power, or for application-specific PSF engineering strategies; other times underfilling is unintentional and due to clipping or limiting apertures in the microscope optical path. In either case, the excitation volume will have a larger spatial extent than predicted by theory for the objective’s listed NA.

Besides the NA and wavelength, other experimental and sample specific factors affect the resolution. For example, the refractive index (RI) varies for materials. Air has an RI close to 1, water has an RI around 1.33, and different types of glass can exhibit RIs in the range from about 1.5 to 1.9. When light rays are converging to a focus, discontinuities or changes in the RI can cause marginal rays to be refracted more strongly than rays close to the optical axis, thus degrading the quality of the focus. This condition is a type of optical aberration called spherical aberration (SA), and SA will reduce the actual spatial resolution from the theoretical prediction. This resolution degradation is typically more pronounced in the axial direction, the direction along which PSFs often have their longest spatial extent. SA can be reduced by matching the RIs of the immersion medium and the sample, when possible. Some microscope objectives correct for some SA through compensation mechanisms such as a correction collar, which can restore diffraction-limited imaging over a range of RI mismatches. In other systems adaptive optics such as deformable mirrors can compensate for such aberrations^42,43^. Although SA could be corrected, RI mismatch also gives rise to a different focal shift for the imaging plane from the travelling distance of the stage (or the objective) along the z axis. This unequal displacement results in axially distorted images, either compressed or elongated. A correction factor can be calculated to correct for this axial distortion and should be applied^44,45^ (**Fig. 7**).

**Figure 7.**
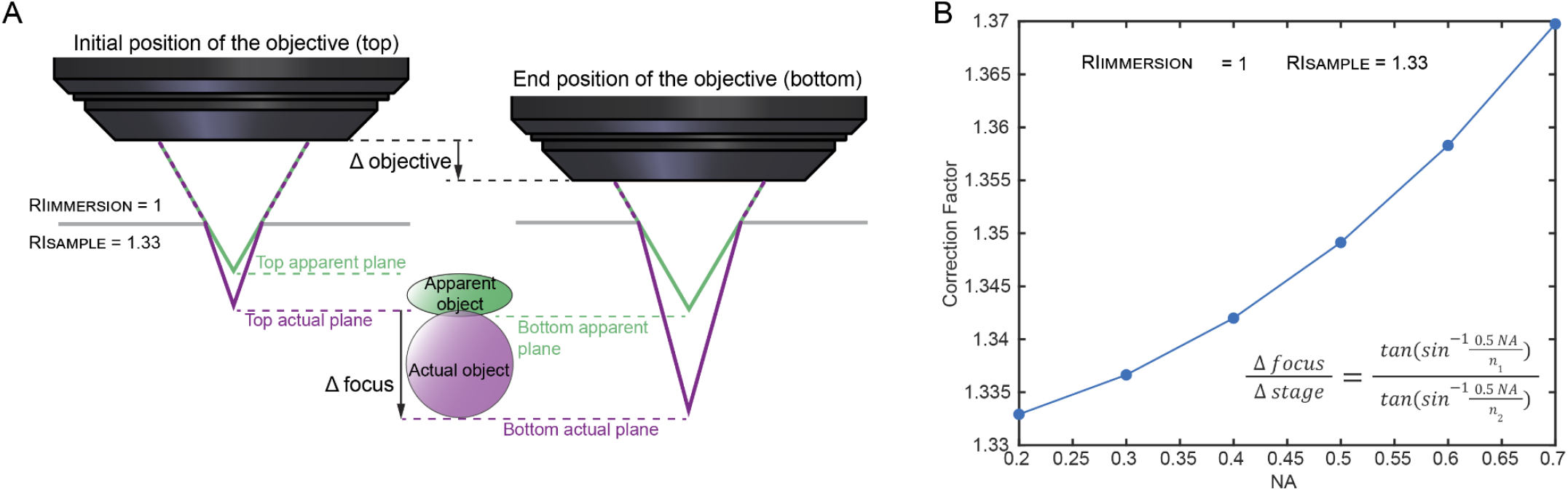
Object distortion caused by refractive index mismatching and the correction factor. (modified from Fig. 1d in the Diel et al.^45^). **A**. An objective lens is scanned axially (black arrow, Δ objective) through a spherical object (‘actual object’, purple) to obtain a 3D image. If there is no index change between the objective and the focal point, there is no refraction occurring (green rays). The distance the objective moves (black arrow, Δobjective) is equivalent to the distance moved by the focal plane (green horizontal lines). However, if the light rays travel from air (RI_IMMERSION_ = 1) into water (RI_SAMPLE_ = 1.33), there is refractive index mismatching, and thus refraction occurs (purple rays). In this condition, the movement of the objective (black arrow, Δobjective) differs from the movement of the actual focal plane (purple dashed lines, Δfocus). As a result, when a spherical object (‘actual object’, purple) is imaged, the rendering of this sphere (‘apparent object’, green) appears compressed in the axial direction because the acquisition software has assigned the objective’s travelling distance (Δobjective), not the focal plane’s travelling distance (Δfocus), to the object’s z axis. (B) A plot shows the correction factor converting the movement of the objective (Δobjective) or the z-stage (Δstage) to that of the actual imaging plane (Δfocus) as a function of NA of the objective. RI_IMMERSION_ = n_1_ = 1, and RI_SAMPLE_ = n_2_ = 1.33.

Overall off-axis aberrations (e.g., coma and astigmatism) intrinsic to the lenses gradually increase and deteriorate the geometry of the excitation volume toward the periphery of the field-of-view. The aberrations are the result from the combination of relay optics, objectives, the arrangement of scanners (closely-coupled or separately-conjugated orthogonal scanners), and so on. Measuring the resolution across the FOV is recommended, especially for the large field-of-view imaging system.

It is worth noting that tissue scattering deteriorates the resolution in a depth-dependent manner, as the marginal rays out of the objective travel a longer distance than an on-axis ray, and accumulate larger scattering and attenuation. The difference in travelling distance can result in a smaller effective NA and impacts the effective excitation volume as the imaging plane goes deeper, especially for high-NA objectives^46,47^. This resultant degradation in signal and resolution can be more severe than simply starting with a lower effective NA, and is a reason why it is generally recommended to underfill objectives when imaging deep^48,49^.

Note that the excitation volume measured in this protocol is focused on characterising the imaging system only. Imaging biological specimens will entail additional aberrations and scattering, which can be compensated for by adaptive optics^50^.

As the resolution is a function of wavelength, the measurements should be performed with the wavelength used in the experiments (**Fig. 6**). Lens designs, antireflective coatings, dichroic filters, and mirrors all have wavelength-dependent properties and can influence the measurements made.

##### Excitation volume measurement

A standard method to measure the excitation volume shape is to image fluorescent beads that are small enough to approximate an “impulse” function. That is, fluorescent beads that are smaller than the expected excitation volume. Beads with a diameter of 0.2 um are commonly available, and are smaller than most multiphoton systems’ excitation volume when near-infrared (>700 nm) light is used. These are sometimes called “sub-resolution” beads. The beads can be spread across a microscope slide or embedded in a 3D tissue phantom (e.g. agar). Of course, even these small beads are not true impulse functions, but in practice they are a close enough approximation that the resulting measurements are useful for characterising a system. Imaging small structures *in vivo* can provide a rough proxy for bead measurements. However, biological features are not proper impulse functions either, and they are typically extended in some dimension. Moreover, a non-zero background signal can distort excitation volume measurements from small structures. Therefore, measurement using sub-resolution fluorescent beads on a slide provides a better baseline for comparisons over different time points and different imaging systems.

### Materials

#### Equipment

1. Agarose (e.g., #16520050 Invitrogen)
2. Eppendorf tube 0.2 μm beads (e.g., F8811 Invitrogen)
3. Glass slide with a concavity, or well, in the centre (e.g., #632200 Carolina)
4. Motorised vertical translation stage with ≤ 0.5 μm minimum incremental movement

#### Equipment setup

1. Make a 0.75% agarose solution. For example, dissolve 0.150g agarose (e.g., #16520050 Invitrogen, low melting point agarose) in 20 ml water (∼75° C), and vortex until completely dissolved. Note that low melting point agarose is recommended for the ease of sample preparation.
2. Transfer 1000 μl of the agarose solution to an Eppendorf tube, and wait for the agarose to cool slightly to ∼ 50° C– just warm to the touch.
3. Add 0.2 μm beads (e.g., F8811 Invitrogen) in 0.75% agarose solution in the ratio of 1 : 1000∼10000 in volume. For example, add 0.2 μl of bead solution to 1000 μl agarose solution in the Eppendorf tube. Vortex the mixture for ∼ 5 seconds and then spin it down (∼5 seconds with a mini-centrifuge) to make the mixture uniform and remove bubbles.
4. Transfer the bead/agarose mixture to a glass slide with a concavity, or well, in the centre (e.g., #632200 Carolina), and cover with a cover glass with a known thickness (e.g., #1, #1.5, …). Completely fill the space between the slide and the cover glass with the mixture, leaving no air gap. Avoid bubble formation during the process. Air gaps and bubbles can move around during the imaging session, and cause the movement of the beads.
5. Wait for ∼ 15 minutes for the mixture to completely cool down and solidify.

OPTIONAL: Seal the sides of the cover glass (with nail polish, wax, optical glue, and so on) to prevent evaporation over time. Evaporation also causes the sample to move. If the measurement is done within ∼ 2 hours, sealing is not usually necessary.

### Procedure

#### Spatial resolution measurement

1. Place the prepared slide on the motorised stage, and search for the beads in the centre of the field-of-view. A field-of-view size of 100-200 μm can be used for an initial survey and finding the imaging plane.
  a. Note that in order to most efficiently perform two-photon excitation, the shortest pulse width in the imaging plan should be used and thus the GDD compensation procedure is suggested to perform before this protocol (see Grouped Delay Dispersion Optimisation).
  b. Inefficient excitation requires the usage of higher power to achieve an acceptable signal-to-noise ratio. The consequence of using higher power would result in photodamage or drifting of the beads during the measurement.
2. Zoom-in to an isolated bead with an acceptable brightness. Make sure you’re imaging a single bead, and not a cluster of beads. This can be done by visually evaluating the average size of beads found in the FoV and choosing one or a few smallest beads. The size of the zoom-in field-of-view could be ∼ 10-30 μm (512 pixels) so that the pixel size is small enough for sufficient sampling, and the field-of-view is large enough to cover the entire bead with enough margin around the bead for correct data fitting.
  a. Note that overly zoomed-in field-of-view may not bring further advantages but instead cause disadvantages. The beads could photobleach quickly and in extreme cases, the excess heat deposited locally could melt the agarose.
  b. Before choosing a bead for measurement in the next step, move the beads in and out of the focal plane to get a sense of what total axial scan range is required, and also check if there are any other beads in close proximity in the axial direction, which would interfere with proper measurements. Examine the brightness and ensure the brightness at any image plane does not saturate the PMT or plateau the dynamic range. If that occurs, readjust the laser power or the gain of the PMT, or use another bead.
3. Acquire a z-stack of the bead with a proper step size and a total range.
  a. Select the z-extend of the stack to be 4-6 times larger than the theoretical axial extent of the PSF. For example, for a system with an axial FWHM of 5 μm in theory, 20-30 μm of total scan range is suggested.
  b. Typically, for more robust fits in the later analysis steps, 10-100 pixels per bead (or 0.02-0.06 μm per pixel) in both XY and 10-100 slices (or 0.1-0.5 μm of step size) in Z are suggested.
  c. At each focal plane, several frames can be acquired and averaged to yield a high signal-to-noise image.
  d. Note again that minimising the power used for the measurement helps avoid thermal and bleaching issues, and operation below the saturation regime results in correct resolution measurements.
  e. It is recommended that beads are measured at different locations across the field-of-view and at different depths, as optical aberrations tend to increase away from the centre of the field-of-view, and the spherical aberration varies and the scattering increases over the imaging depth.
  f. Beads at the edge of the field-of-view can be accessed by offsetting the scan centre of the linear galvanometer scanner. For example, systems equipped with a x-resonant scanner and a y-linear scanner can shift the centre of the zoom-in field-of-view along the y axis. For systems equipped with a x-linear scanner and a y-linear scanner, the centre of the zoom-in field-of-view can be positioned at any point throughout the accessible field-of-view. This offsetting function might appear ‘shift’, ‘offset’, or ‘park’ on the microscope control panel.
4. Fit the intensity measurement with Gaussian curves in the lateral (XY) and axial (Z) direction, respectively (**Fig. 8**). The FWHM reported, in units of length (e.g., nm or um). GitHub - SWC-Advanced-Microscopy/measurePSF: Measure PSF FWHM along different axes. In cases where the excitation volume is tilted in Z, the Gaussian fit procedure should account for this tilt, otherwise the excitation volume will be underestimated. The current version of our example code does not include a subroutine to correct the tilt. One could use the built-in rotation function, such as ImageJ (‘Image->Transform->Rotate…’) and MATLAB (‘imrotate’), to correct the tilt.
  a. Note that if an air immersion objective is used, due to the difference between the refractive index of the objective immersion medium (e.g., air) and the specimen medium (e.g., water), the actual focal position (Δfocus) within the specimen was moved a different amount from the stage movement (Δstage). Therefore, a correction factor is required to convert the axial stage movement to the actual focal movement. According to the reference^45^, a simplified calculation of the correction factor is

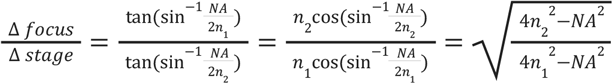

where n_1_ is the refractive index of the immersion medium, n_2_ is the refractive index of the sample, and NA stands for numerical aperture of the objective.
5. For full characterization, the measurement should be performed several times, for many locations in the field-of-view: the centre, the edges of the X scan, the edges of the Y scan, and different depths. It is useful to show where resolution breaks down. Instead of simply reporting the best values, show where the resolution starts to degrade and by how much. For routine checks, measuring at just two or three reference locations can be sufficient.

**Figure 8.**
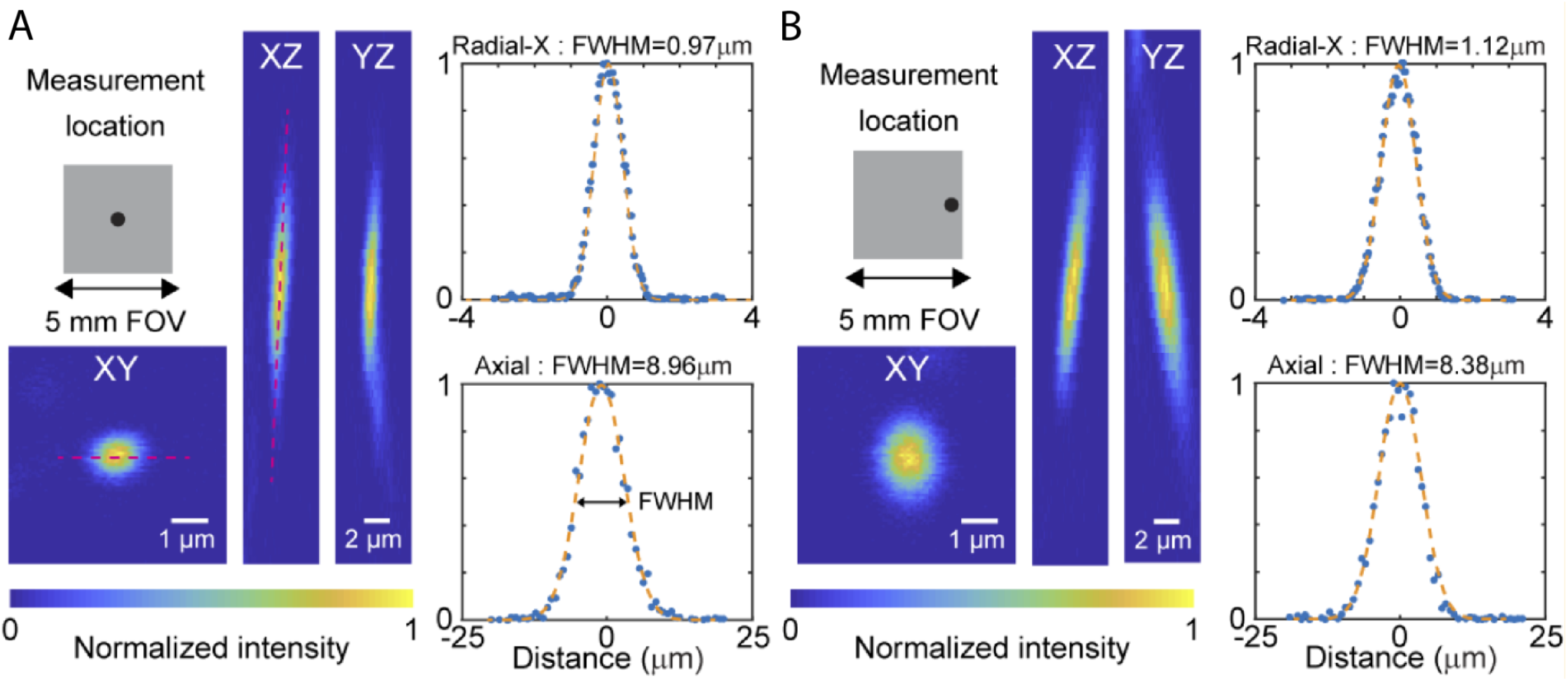
Example measurement of PSFs. 0.2 μm fluorescent beads were embedded in 0.75% agarose gel. 40 μm z-stacks were acquired at the depth of 500 μm. Beads at the centre **A** and the edge **B** of the field-of-view were measured. The example images are shown from the XY, XZ, and YZ cross-sections, respectively. The intensity profiles of the beads (red lines) in the X direction on the XY plane and in the Z direction are plotted, which are fitted to a Gaussian curve (orange dashed line) to extract the radial-X and axial FWHM of the PSF. Note PSF degradation for lateral position of the bead. While small degradation of resolution at the edges of FoV is often expected by design, anything beyond that is usually caused by misalignment in the optical path and should be revisited and corrected.

### Troubleshooting

Individual beads should be isolated from other beads to ensure correct estimation of the excitation volume geometry. Do not necessarily search for the brightest points because they might be clumps of more than one bead.

Generally, the laser power used should be close to the minimum required to clearly identify individual fluorescent beads. If higher powers are used, there is a risk of over- or underestimating the excitation volume, through either saturation or bleaching respectively, and this issue is exacerbated when using fluorescence molecules with relatively large cross sections^31^.

When referring to a theoretical or expected resolution, be sure to include the equation, the values for any parameters in the equation, and a paper reference, e.g. Zipfel et al. 2003^31^. Be clear whether FWHM, 1/e radius, or 1/e^2^ radius of the PSF is reported. This provides clarity to the reader and enables apples-to-apples comparisons.

The lateral pixel size needs to be carefully calibrated. Do not simply take the numbers off of the manipulator controller unless you are certain it is calibrated correctly. Calibration can be performed with a structured sample (e.g., a grid) with known dimensions (see Field of View Size).

The axial movement of the stage needs to be calibrated, as well. Again, do not blindly rely on the numbers given by the manipulator controller unless it is reliably calibrated.

Tissue phantoms^51^ can provide a more realistic measurement environment (compared to glass slides or agar blocks). However, as these are not standardised, it can be difficult to compare results from different labs using different phantoms.

### Pulse Width Control and Optimisation

#### Introduction

Nearly all multiphoton microscopy makes use of modelocked^52^ ultrafast lasers with pulse durations on the order of 100 femtoseconds (100 × 10^−15^ seconds). Due to the non-linearity of multi-photon excitation, peak intensity matters more than average power for efficient excitation. The efficiency of multi-photon excitation using pulsed lasers versus CW (continuous wave) lasers, with the same time-averaged power, is given by:

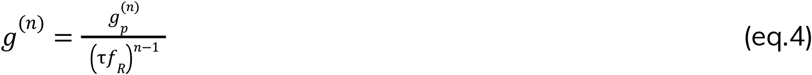

where *g*^(n)^ = enhancement factor, τ = pulse width, *f*_*R*_= repetition rate, n = number of photons in the absorption process^32^. The factor 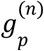depends only on the pulse shape, and is equal to one for a rectangular pulse and 0.59 for a hyperbolic secant (sech^2^) envelope, which is close to the typical shape of the pulses delivered from modelocked pulsed lasers^52^. For two-photon imaging, with a standard Ti-Sapphire laser operating at 80 MHz with 150 femtosecond pulses, this enhancement is ∼ 50,000, and is strongly dependent on τ, the pulse width.

The generation of short laser pulses requires finite bandwidth - the shorter the desired laser pulse, the broader the spectral content. This can be rigorously derived classically through Fourier transform relationships, and for the sech^2^ envelope mentioned above, leads to τ _*p*_ ×Δ*v* _*p*_ ≥ 0. 3148, with τ _*p*_ the full-width at half-maximum (FWHM) of the laser pulse envelope, and Δ*v*_*p*_ the FWHM of the frequency spectrum ^52^. Converting frequency to wavelength means a 150 fs pulse centred at 920 nm needs at least a FWHM bandwidth of 6 nm, a 100 fs pulse 9 nm, and a 50 fs pulse 18 nm. It is important to note that broad bandwidth alone does not *guarantee* a short pulse, but is *required* to generate one. When the pulse is the shortest it can possibly be, given the nominal bandwidth envelope, the pulses are known as *transform-limited pulses*.

Once the light pulse leaves the laser, it can be broadened in time through interactions with dispersive materials, like glass, resulting in longer pulse duration. A dispersive material is one that has a frequency (wavelength) dependent refractive index. If the refractive index increases with increasing frequencies (decreasing wavelength) the material is said to have positive dispersion, whereas if the refractive index decreases with increasing frequency it is said to have negative dispersion. At the common wavelengths used in microscopy, most materials exhibit positive dispersion^53^. The refractive index can be related to the speed at which light travels through a medium; for positively dispersive media, the bluer portion of the pulse travels at a slower velocity than the redder portion, temporally broadening or “stretching” the pulse, and resulting in a “chirped” pulse. The more material the pulse propagates through, the greater the temporal broadening or stretching. Microscopes with elaborate optical systems (such as large FoV systems^28^) have more or thicker glass elements and so produce more dispersion. Similarly, head-mounted multi-photon systems that require a long fibre optic cable for beam delivery will generate a lot of dispersion (unless specialised fibres are used)^54,55^.

The broadening effect can be counteracted with a process called dispersion compensation, where the user purposefully introduces a fixed magnitude of dispersion that exactly matches that of the microscope’s optical path, but of the opposite sign, such that the combined dispersion from the compensation unit and microscope sums to zero net dispersion. Some lasers have dispersion compensation included as an integral component option (e.g. Coherent Vision and Axon, Spectra Physics Mai-Tai DeepSee and others). For lasers that do not include dispersion compensation, either an external commercial or home-built dispersion compensation unit is used, or in many cases, no compensation system is present, and the user has limited opportunity to modify the pulse width, but should still consider the role dispersion may play in their system.

For two-photon microscopy, the main consequence of temporally stretched pulses is that the effective two-photon excitation efficiency drops (see eq 4). The drop in efficiency for non-linear processes is often significant, even though the total average power being delivered to the sample remains the same. For higher order processes, such as three-photon excitation, the fall-off in signal with longer pulse widths is even more rapid. For shorter pulses (<=50 fs) it is especially important to note that the complete “dispersion” relationship also includes higher order terms that may need to be controlled as well^9,56^.

##### Group delay dispersion (GDD) optimization

The art and practice of dispersion control is rich, and strongly dependent on the equipment at hand^57–59^. Beyond that, there is a plethora of instantiations for building external compensation modules, or general dispersion control, the details of each is beyond the scope of this protocol, especially if higher order dispersion control is necessary.

This protocol will assume that the user has a form of an easily tunable dispersion control module, and will use the level of fluorescence generated following two-photon excitation as a surrogate for measuring changes in the pulse width. The goal is to collect image intensity data throughout the range of GDD correction values, and ideally, find the dispersion compensation setting that corresponds to the brightest image. This would correspond to the shortest laser pulse that your system is capable of delivering, and hence the highest peak intensity of the laser, for a given average laser power. The fluorescence can be used because the signal is proportional to the laser intensity squared, while the intensity is proportional to the average power/pulse width, as described earlier. If the lab has invested in an autocorrelator, using that will provide a more quantitative assessment of the pulse width, though the method described above is a reasonable alternative for fast optimization.

If you do not have either dispersion control nor an autocorrelator, how concerned should you be about laser pulse management? Dispersion management/compensation becomes less important for simpler optical paths and longer initial pulses (due to pulse broadening being non linearly proportional to the pulse width). If you have a relatively simple microscope, with few optical elements (e.g. mirrors, galvanometer-based mirror scanners, scan lens, tube lens, objective) between your laser and the sample, and your laser has initial pulses of ∼ >150 fs, dispersion compensation may not be a worthwhile investment. If neither of these conditions hold, it may be worthwhile to consider working with either another lab or a vendor to measure the laser pulse on sample with an autocorrelator to see if additional measures are needed. This is especially important for short pulse laser systems. For example, for a system with a total GDD of 6000 fs^2^, a 60 fs pulsed laser will be broadened to 284 fs, an almost 5x increase, and concomitant ∼ 25x loss of two photon efficiency, while a 150 fs laser pulse will be broadened to 186 fs, only a 1.24x in pulse duration, with a two-photon efficiency loss of only 1.54x.

### Materials

#### Equipment

- Stable fluorescent sample slide e.g. pollen grain slide (Mixed pollen grain slide, Carolina Biological Supply Company), fluorescent acrylic slide (Chroma, etc.), fluorescein bath (for water immersion objectives) or a fluorescein drop on a microscope slide under a coverslip.
- Microscope control software that allows for live image histogram viewing.
- Objective lens used for imaging

#### Equipment setup

1. Turn on the laser and allow the microscope to stabilise (usually between 30 minutes and 1 hour)
2. Set the laser wavelength to one used for imaging experiments
3. Start the microscope control/image acquisition software
4. Install a microscope objective lens used for imaging experiments
5. Place the sample under the objective lens
6. Apply any immersion media appropriate for the objective lens and position the lens at roughly its focal distance away from the sample

### Procedure

1. Begin live scanning and focus on the sample.
2. OPTIONAL: If using a slide with features (e.g. pollen grains), search for the features in the sample using the stage to navigate. NOTE: The laser power may need to be adjusted to find features more easily. CAUTION: Relatively high laser power can easily burn pollen grains, so reducing the laser power may be necessary.
3. OPTIONAL: If using a slide with features (e.g. pollen grains), use a higher magnification so that the smallest feature chosen occupies ∼ 50 × 50 pixels. If sized like this, it ensures that any small motion or pixel alignment errors will not significantly corrupt the measurements. However, if magnification is too high, bleaching and damage to the pollen grain may occur.
4. Open a live histogram of pixel intensity values for the image.
5. Set the vertical scale (number of pixels) to “logarithmic” to see the low pixel counts such as those between 10^0^ and 10^1^.
6. Set the x-axis to show the full range of possible pixel values and adjust laser power and/or PMT settings so that live pixel values occupy 25% of the full range. Once these laser power and PMT settings are set, these cannot change for the duration of the measurements. If they change, the steps below will need to be repeated. NOTE: if the signal level is very high (values near the maximum), the preference would be to first turn down the laser power, rather than the PMT voltage. Lower laser powers result in less possible photobleaching, which would interfere with the interpretation of the measurement.
7. Adjust the pixel intensity range on the histogram to show 50% of the full range (see **Fig. 9**), and record/remember the maximum pixel value. If the histogram display also shows the mean and max values of the image, record that too.
8. Continue imaging for at least 2 minutes, and compare the current pixel values with the values noted in step 5. If the values are significantly different, it shows that either the sample is bleaching and/or the system is not stable. Find either a more stable sample, or turn the laser power down to decrease photobleaching, such that the pixel histogram is stable for ∼ 2 mins. CRITICAL: The signal needs to be stable to continue this protocol because the absolute pixel values need to be compared across time as the dispersion is adjusted.
9. Record the current dispersion compensation setting of the laser (usually in fs^2^ GDD) or external module and record the shape of the histogram alongside the mean and max pixel values.
10. 1Change the dispersion compensation by a fixed amount (increments of ∼ 2000 fs^2^ GDD is generally sufficient).
11. Examine the live histogram and again record the shape of the histogram alongside the mean and max pixel values.
12. Continue changing dispersion compensation in both the positive and negative directions from the initial setting. If the mean pixel value drops >30% from that of the initial recorded value, you do not need to continue in that direction.
13. Plot mean pixel values of the image as a function of dispersion compensation values (**Fig. 10**). Identify the GDD value that gives the highest mean pixel value.
14. Now repeat steps 9 to 13, but for GDD compensation increments of ∼ 250 fs^2^ GDD. The dispersion compensation setting that gives the highest mean pixel value is the one you want to use for your experiments.
15. Repeat steps 6 to 14 for different wavelengths and objectives that will be used for imaging.

**Figure 9.**
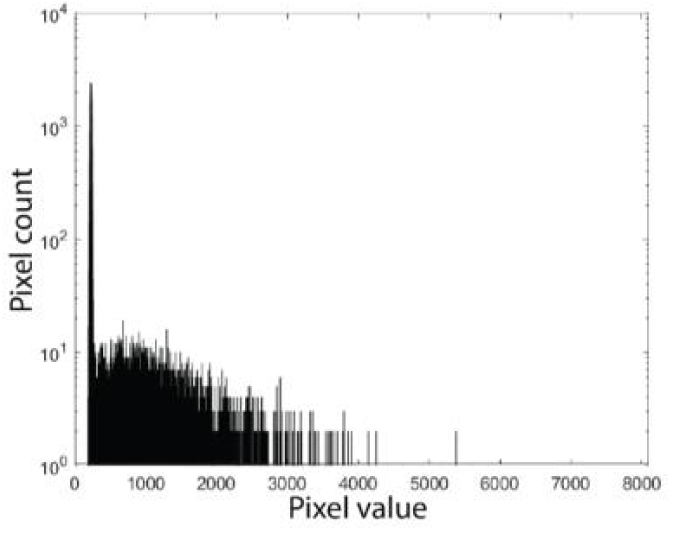
Example pixel histogram used to optimise laser pulse width.

**Figure 10.**
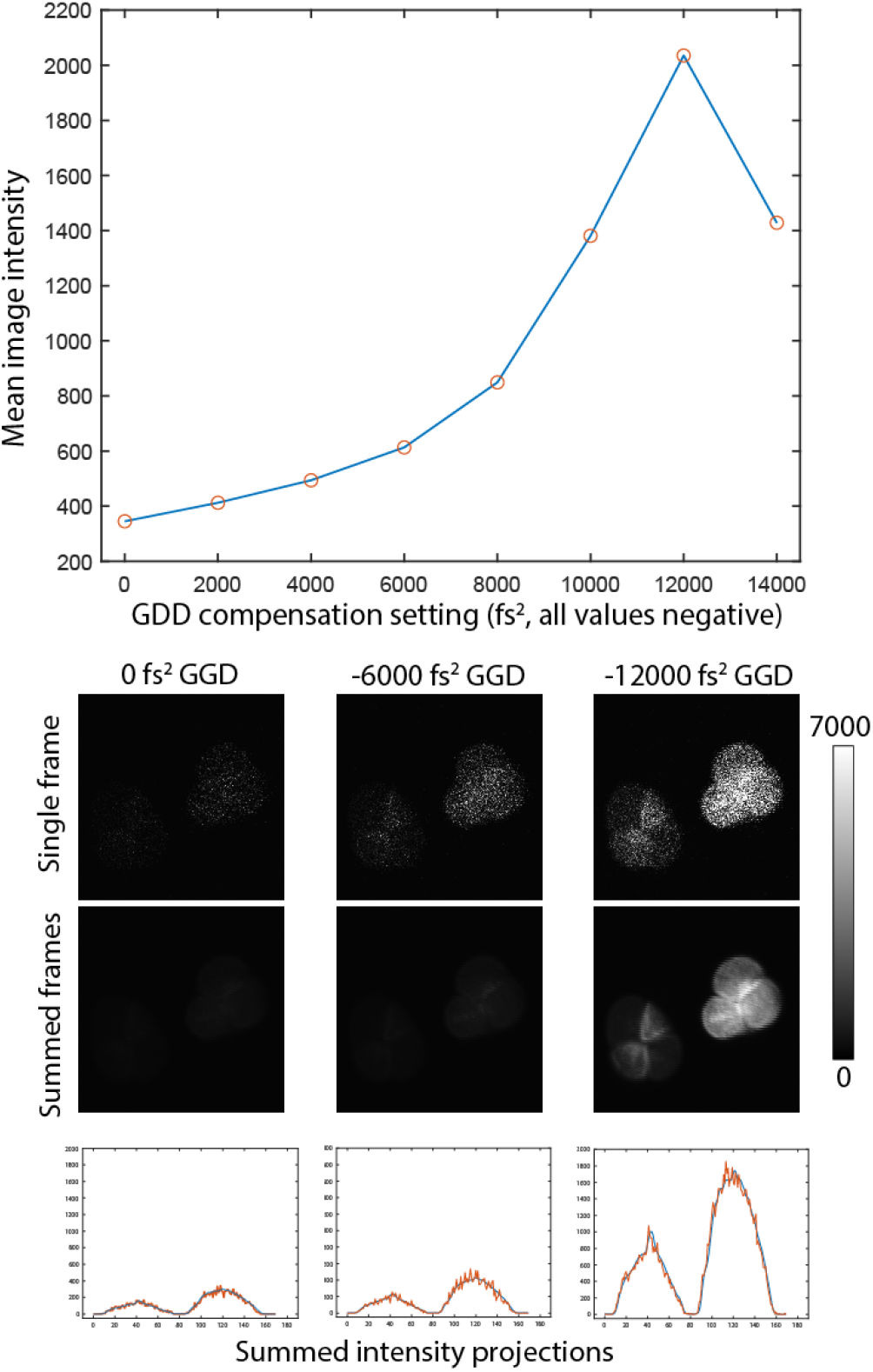
Pulse width optimization measurement. **A**. Plot of mean image intensity vs GDD compensation value for fixed average power. For this system ∼-12000 fs^2^ of compensation is necessary for the microscope to achieve highest excitation efficiency. **B**. Example images acquired of the different settings of GDD compensator, and projected intensity plots that can be used for analysis described in the section.

### Troubleshooting

It may be the case that the signal intensity never declines in one direction, but instead, continues to rise until the limit of your dispersion compensation correction is reached (this happens occasionally for internal correction modules). When this happens, the user has to either decide to accept this value as the “best” correction, or must consider adding more negative dispersion through an additional external unit. If the user typically uses a fixed wavelength, multiple bounces between chirped mirrors provides a straightforward way to add dispersion^60^.

One interesting aside about mirrors, is the complex coatings of dielectric mirrors can have very strong and very specific wavelength dependence of dispersion (**Fig. 11**). Thus despite the high reflection efficiencies of multilayer coatings, the authors recommend using metallic mirrors if possible, or dielectric mirrors that are designed and calibrated for known amounts of dispersion. If your system has “normal” dielectric mirrors, we recommend the user “scans” the laser by 2-3 nm increments in a 60 nm range around the expected use wavelength, and records the fluorescence signal. If the resulting fluorescent intensity measurement displays any sharp peakiness, it is possible that the dielectric mirrors are significantly distorting the pulse shape. As the total effect of dielectric coatings (particularly old ones) on pulse shape can not be assessed without sophisticated instrumentation, we suggest replacing dielectric mirrors with metal-coated alternatives rather than simply “correcting” for GDD peakiness with a pre-chirper.^61^

**Figure 11.**
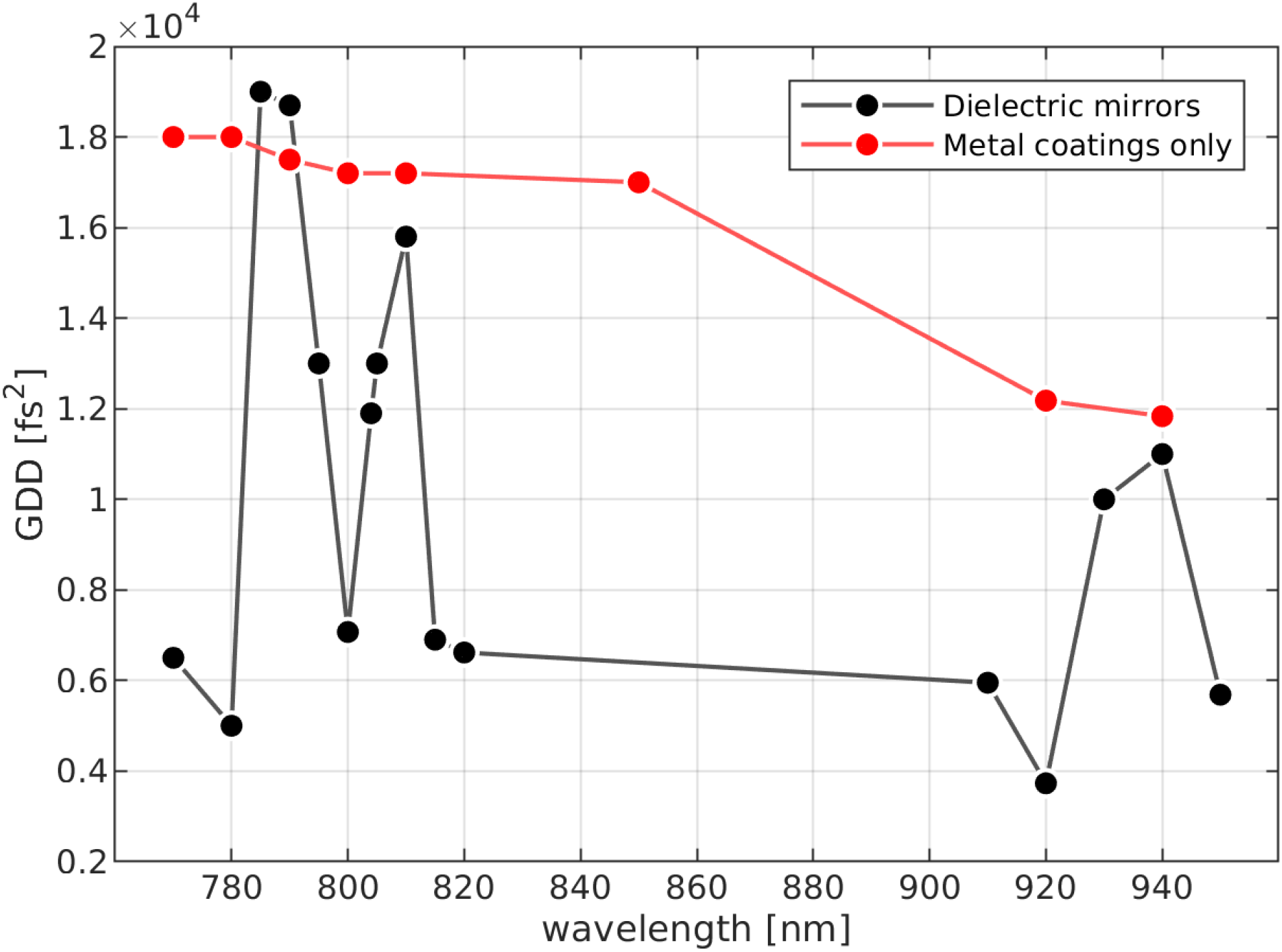
The effect of dielectric coatings on GDD. Optimal GDD was estimated using a fluorescent slide and the built-in GDD compensation on a Coherent Chameleon Vision II laser. The compensation curve was first measured with three dielectric coated (ThorLabs EO3) mirrors in the path (black line). Very obvious sharp peakiness is seen in the required compensation value as a function of wavelength. After these three mirrors were swapped with metallic coated mirrors, the peakiness completely disappears (red line). The three EO3 mirrors did not contribute equally to the above effect (data not shown).

### Photomultiplier Tube Performance

#### Introduction

Most multiphoton microscopes use one or more photomultiplier tubes (PMTs) to detect light. In brief, the objective and collection optics direct photons emitted from the sample to the primary photosensitive element of the PMT, its photocathode. Photons having sufficient energy will cause a photoelectron to be generated via the photoelectric effect and then amplified through a cascade of dynodes (involving a high potential difference distributed across a chain of increasingly positive dynodes). This converts the small number of photoelectrons to a much larger number of electrons at the PMT anode (gain of 10^5^-10^6^). The bolus of charge arrives at the anode over ∼ 5-10 ns yielding a small burst of current. Typically, a transimpedance amplifier (TIA) is then used to convert this photocurrent to an analog voltage and amplify that further. That voltage signal is then digitised by a high-speed ADC card in the acquisition system.

It is important to acknowledge that PMT performance degrades over time. Therefore, the PMT should be regarded as “consumable” and be replaced periodically if the microscope’s detection performance is to be maintained. Most often, degradation is related to the total charge that has passed through the PMT, and this leads to a loss of both cathode sensitivity and photocurrent amplification but can also manifest as an increase in dark current and a decrease in dielectric resistance^62^. Moreover, “day one” performance, as well as rate of decline, vary substantially across units of the same PMT model and the rate of deterioration is influenced by usage (for example light exposure and average anode current).

A quantitative means to characterise PMT performance is therefore useful to (1) select between different units when installing a new PMT into the microscope, (2) diagnose issues that might arise with image quality and (3) benchmark performance over time to make informed decisions about when to replace a PMT.

We note that a lab’s specific performance requirements and financial considerations will also play into decisions around when to replace PMTs. However, by quantifying performance, a consistent policy can be adopted that, alongside other routine tests and maintenance described in this article, should allow minimum standards of data acquisition to be maintained.

In this section, we describe a simple protocol to quantify light detection performance. With minor variations, the procedure can be used to test either the entire collection system (defined here as objective, collection optics and PMT), or test a PMT directly. Direct tests of the PMT require it to be disconnected from the microscope, but otherwise require minimal modification to the protocol. We note that this protocol does not provide measurements in physical units, nor does it test different aspects of PMT performance as might be reported on the datasheet (e.g. anode luminous sensitivity), which requires additional sophisticated instrumentation and is beyond the means of most research labs. Instead, we present a protocol that provides a robust, relative measure of performance that can be used comparatively, e.g to compare several PMT units of the same model or monitor performance of the same unit over time. Our measure is based on ideas from signal detection theory and is not specific to PMTs – indeed, the protocol can be applied to any light detector (e.g. hybrid detectors). Finally, it also allows the user to select the optimal operating point (typically referred to as “gain”) for a specific PMT unit. The gain of a PMT is adjusted by changing the high voltage (HV, typically 500 – 1500 V) that is distributed across its electrodes. In most instances, the user will control HV via a lower voltage control signal (e.g. 0–5 V) that is provided to the socket assembly or PMT module. Users should be aware of how gain is controlled on their specific system. For routine monitoring of PMT performance, we recommend performing this protocol at least once every 6 months.

### Materials

#### Equipment

This protocol requires a light source with a stable output. Such sources are available commercially, e.g. based on closed-loop control of an LED, or can easily be assembled using a tritium vial as an approximately constant light emitter^63^, as we have chosen to do for this protocol. Tritium is a radioactive isotope of hydrogen with a half-life of 12.3 years. Tritium vials contain a small amount of tritium gas and a phosphor coating that produces light by radioluminescence when bombarded with beta particles. Vials are widely available with phosphor coatings that produce various colours including ‘red’ and ‘green’. These work well over the duration of most experiments, although additional care must be taken when analysing measurements made across years; first, the half-life of tritium will cause a well defined decrease in intensity, and phosphor degradation will further impact long term stability^64^.

Components for a tritium light source (OPTIONAL):

- Tritium vial 3×11 mm (e.g. https://edcgear.co.uk/products/tritium-vial-1-5mm-x-6mm-capsule or https://www.mixglo.com/store/c2/Vials.html)
- Epoxy resin
- SM1 lens tube, 0.5 inch (Thorlabs SM1L05)
- SM1 end cap (Thorlabs SM1CP2M)
- Mounted Pinhole, 500 μm (Thorlabs P500D)
- OPTIONAL: Neutral density (ND) filter (e.g. Thorlabs NE10B-A)

#### Equipment setup

##### Assembling a tritium light source (OPTIONAL)

1. Select a tritium vial of the appropriate colour for the detection channel being tested (e.g. ‘red’ or ‘green’).
2. Use a small drop of epoxy resin to affix the tritium vial to the centre of the inside surface of the SM1 end cap and allow the resin to set overnight.
3. Screw the end cap, with the mounted tritium vial, onto the lens tube.
4. OPTIONAL: A neutral density filter can be added inside the tube to reduce light intensity if required (see below)
5. At the open end, attach a mounted pinhole.
6. Label the assembled tritium source with a unique reference number, assembly date and ‘colour’ of the tritium vial (red, green).
7. Once assembled, the tritium source can be used for many years. To ensure stability, it must not be assembled/disassembled and should be stored in a dust-free container (**Fig. 12**).

**Figure 12.**
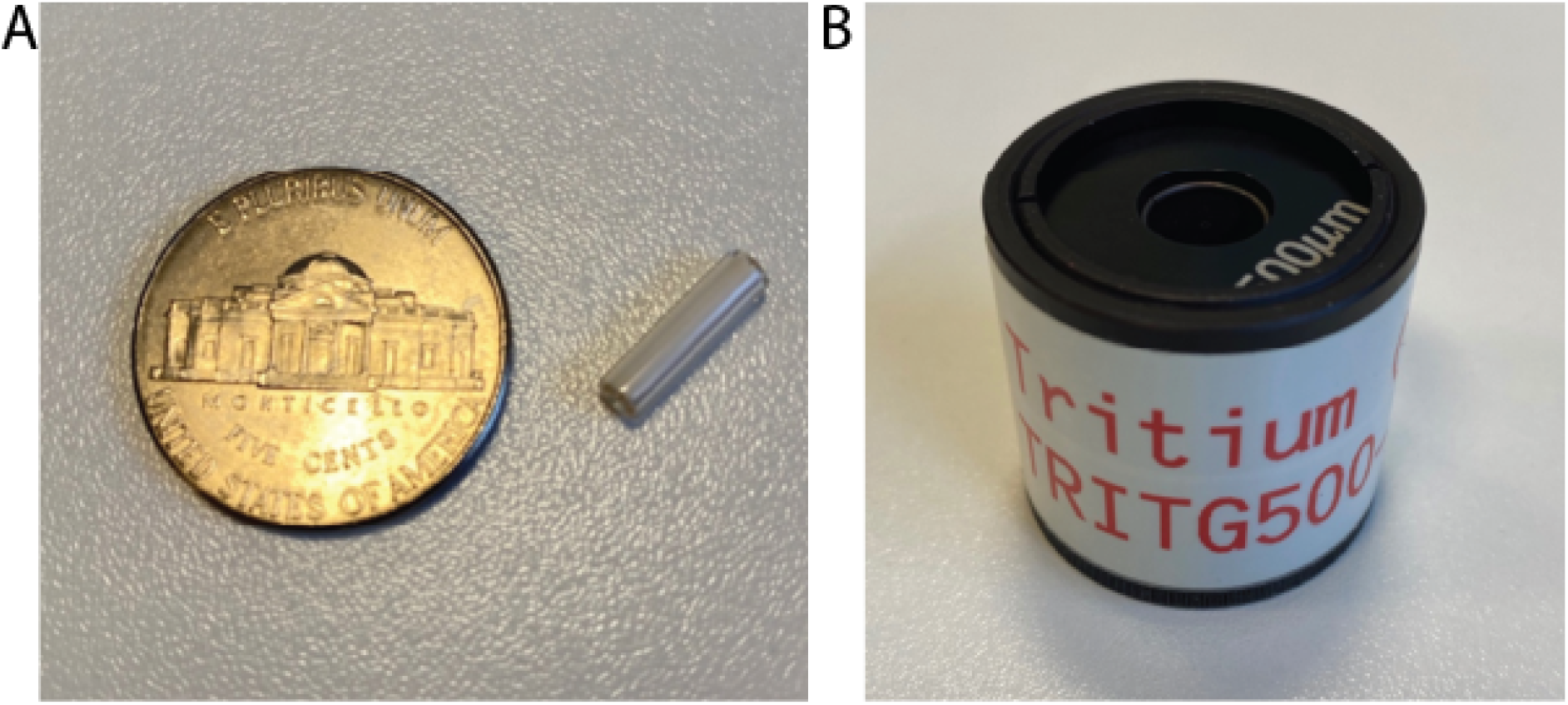
A hand-made tritium light source. **A**. 3 × 11 mm tritium vial next to a 5 cents coin. **B**. The assembled tritium light source. The pinhole is at the top and will be placed immediately beneath the objective to test the entire collection system, or PMT window to directly test the PMT.

##### System setup

1. PMT performance is dependent on temperature. Therefore, room temperature should match that at which experiments are routinely conducted and be consistent within and across tests.
2. For testing the entire collection system, the PMT must be installed in the microscope. In this case, take care to select a light source of appropriate colour (e.g. a ‘red’ tritium light source) according to the detection channel being tested. For testing a PMT in isolation, the PMT will not be installed in the microscope but rather should be mounted within custom optomechanics such that the photocathode can be placed at a fixed distance from the light source. Again, we recommend using a light source whose nominal colour corresponds to the intended use of the PMT.
3. Condition the PMT by operating it at a normal gain(voltage) while shielding it from any light for around one hour prior to testing to ensure stable operation. For new PMTs, or a PMT that has been unused for several months, it may be desirable to first “age” the tube by operating it for several hours as this can improve stability [1].
4. Refer to previous measurement metadata, where relevant, to ensure consistency of instrument settings across tests. Relevant factors include transimpedance amplifier settings, microscope hardware (including emission filters), and image acquisition settings.
5. Precautions should be taken to prevent any stray light from reaching the PMT, such as a light-tight chamber surrounding the microscope and room lights being switched off.
6. Lasers should be switched off, or shuttered, to prevent any laser light from affecting measurements of PMT performance.

### Procedure

#### Collect dark response image series

First, it is necessary to measure the response of the detector under dark conditions.

1. Configure the microscope image acquisition software to collect image data with settings such as 512 × 512 pixels under normal scan parameters, with 16-bit image depth and .tif file format. NOTE: Do not implement any (line or frame) averaging.
2. Collect a short time-series of images (few seconds duration) at each PMT gain setting for a range of gain values. A time-series is taken such that a middle-of-series single image can be used for analysis, and to allow for a quick inspection of consistency, and avoidance of artefacts on the “start” or “stop” frame. For example, for testing the Hamamatsu R10699 with C6270 socket assembly, we used a range of control voltages from 0 to 3.5 V, in 0.25 V steps, which corresponds to 0–900 V across the PMT electrodes. NOTE: An image must be acquired at zero gain to determine any image value offsets unrelated to the PMT itself, and care should also be taken not to exceed the maximum HV indicated on the datasheet for the specific model of PMT.

#### Collect light response image series

Next, the response of the detector to the constant light source must be measured.

1. Place the constant light source in front of the objective (if testing the entire collection system) or PMT window (for direct PMT tests). If using the tritium light source described above, place the pinhole directly beneath the objective. Move the objective very close to the pinhole, for example to the lip of the pinhole mounting. Do not use immersion media.
2. Begin streaming live image data from the PMT. CAUTION: If this is a new tritium light source, or a new model of PMT, take care to slowly increase the PMT gain towards the usual operating range. Image brightness (mean pixel grayscale value) should be similar to biological samples typically used in the lab. If the images appear too bright, add a neutral density filter to the light source (see above).
3. Adjust the lateral position of the constant light source using the X/Y stage controls to maximise image brightness and thereby ensure the pinhole is directly beneath the objective.
4. Collect a series of images at the same range of PMT gain settings that were used for the dark response tests.

#### Data Processing and Analysis

We will illustrate the data analysis and interpretation using:

1. Measurement data from several units of the same multialkali PMT model (Hamamatsu R10699) when first installed (“day one” performance) (**Fig. 13**).
2. Measurement data from a single R10699 when first installed and then after a long period of routine use (**Fig. 14 & Fig. 15**).
3. Measurement data from two GaAsP PMT units (Hamamatsu H10770PB-40) (**Fig. 16**).

**Figure 13.**
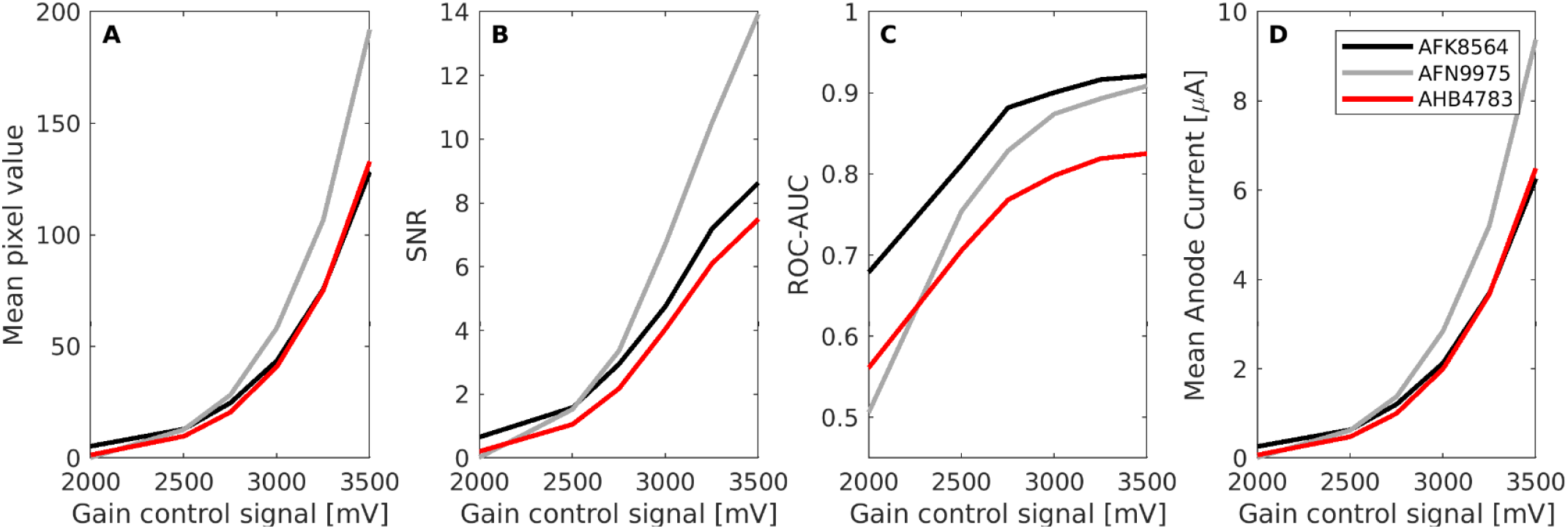
First-day performance for three multialkali PMT units of the same model. Mean pixel value (**A**), SNR (**B**), ROC-AUC (**C**) and mean Anode Current (**D**) data shown for three Hamamatsu R10699 PMTs on the first day of installation. PMTs were tested within the full collection optics system (green channel) of the same microscope. Gain setting is represented by the control signal voltage.

**Figure 14.**
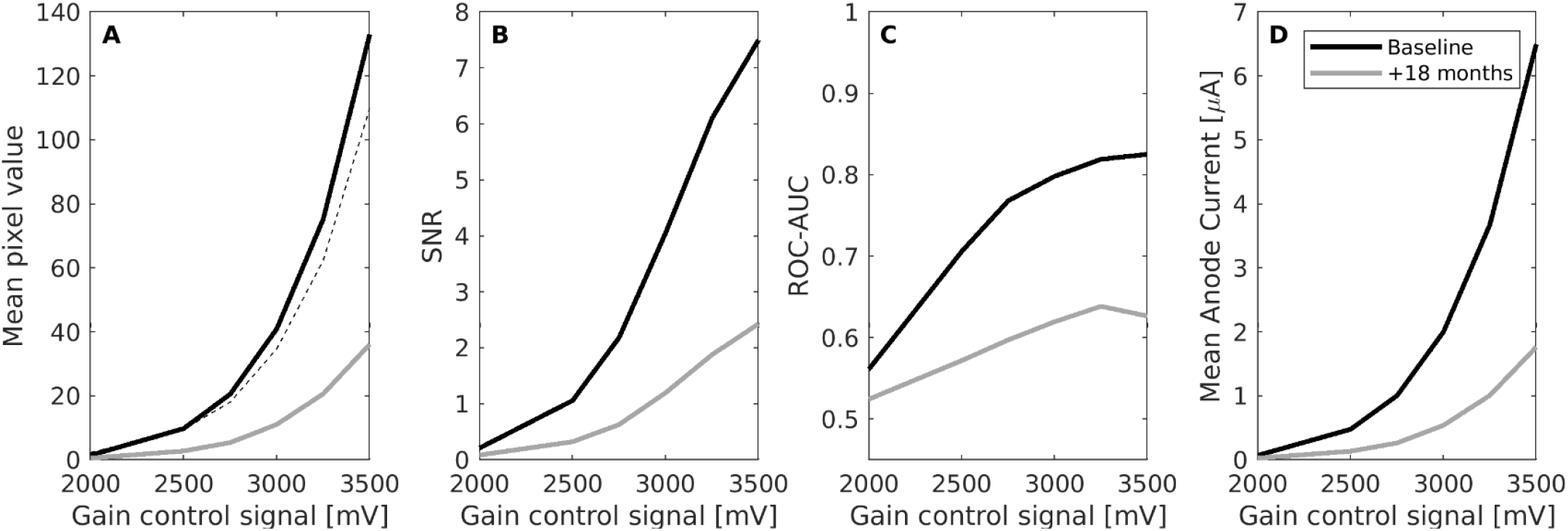
Change in PMT performance over time. Mean pixel value (**A**), SNR (**B**), ROC-AUC (**C**) and mean Anode Current (**D**) collected for a Hamamatsu R10699 PMT unit when it was first installed and after 1.5 years of routine use. Dashed line shows pixel values expected based upon tritium decay alone.

**Figure 15.**
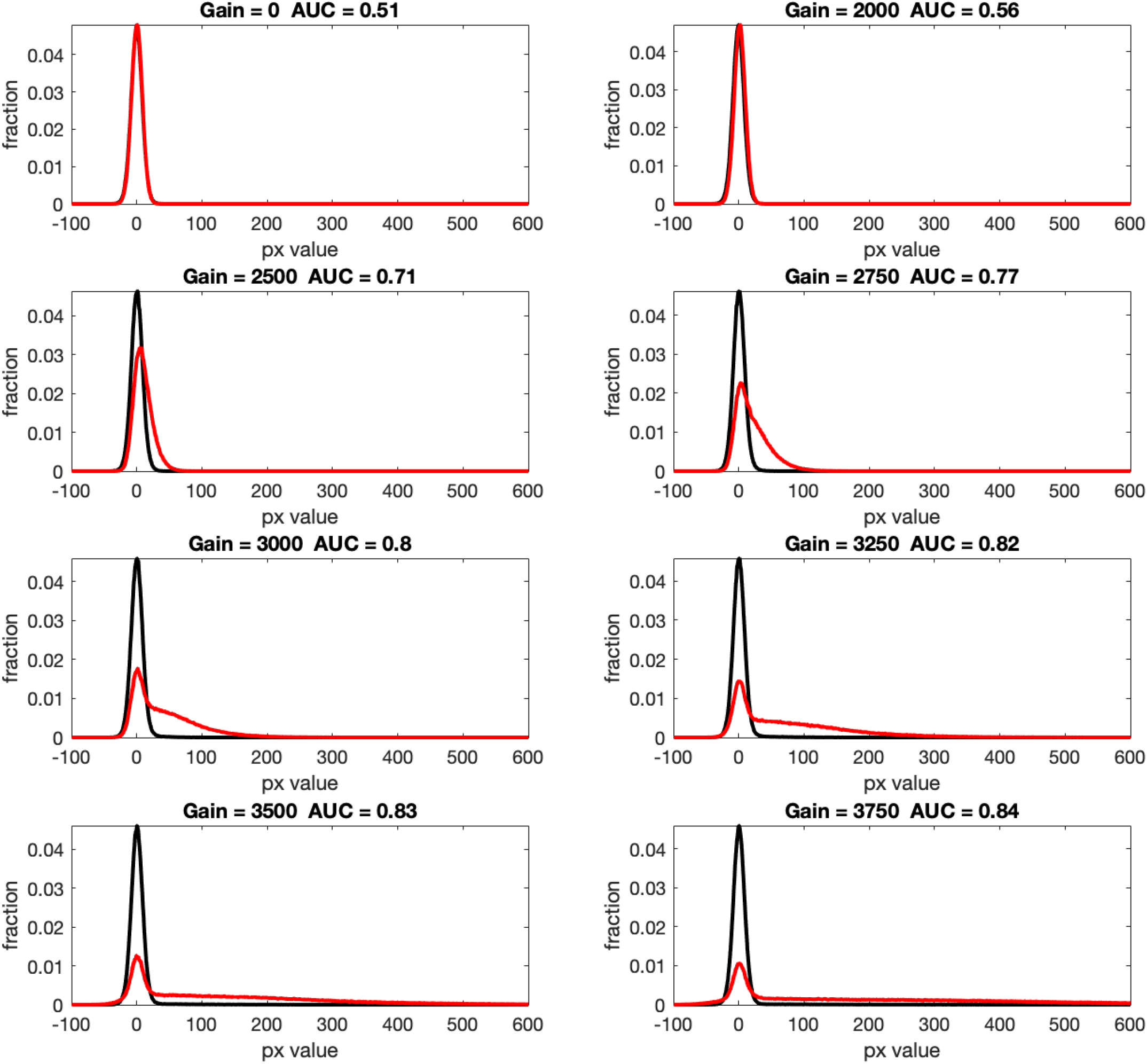
ROC-AUC analysis for an example multialkali PMT. For each gain setting, distributions of pixel values from dark response (black) and light response (red) images are shown and corresponding ROC-AUC values are indicated. Gain expressed as control signal voltage [mV].

**Figure 16.**
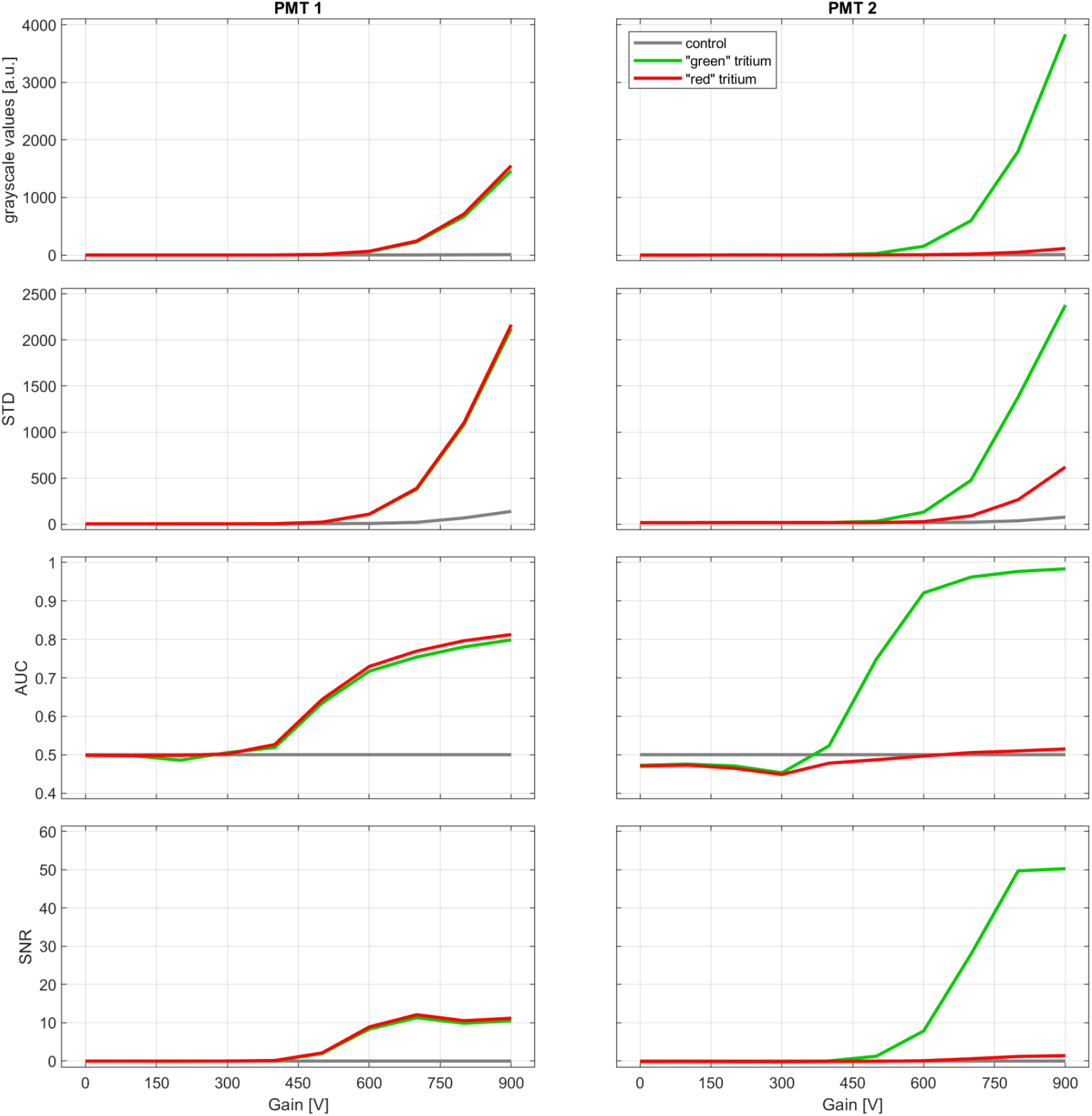
Comparison of two different GaAsP PMTs of the same model (Hamamatsu H10770PB-40). Mean (row 1), standard deviation (row 2), ROC-AUC (row 3) and SNR (row 4) for two example GaAsP PMTs for two different tritium light sources (red & green) and control (no light source). PMTs were measured with bandpass filters in place (PMT1: 570-620 nm bandpass; PMT2: 500-550 nm bandpass). Gain expressed as low voltage control signal [mV].

#### Image processing

First, some basic image processing is required.

1. Calculate the ‘black point’ by computing the mean pixel value in the dark response image acquired at zero gain.
2. Subtract the black point value from all of the images (both the dark response series and light response series).
3. For each image, compute the mean and standard deviation, across pixels.

#### Pixel grayscale value vs gain

Plot the mean pixel value versus gain (represented as either control signal voltage or HV) for the light response image series.

It should be observed that the mean pixel value increases as a function of gain, owing to increased PMT amplification. In Figure 13, which compares “day one” performance of three R10699 units, AFN9975 produces the highest pixel values as expected from it having the highest test sheet anode luminous sensitivity.

Figure 14 shows measurement data for a single unit (AHB4783 from **Fig. 13**) when first installed and then after approximately 18 months of routine use in one author’s lab. Mean pixel values have declined substantially over this time..

Note that these measurements were taken using the same tritium light source. Therefore, some decline in mean pixel value should be expected due to the radioactive decay of tritium. This can easily be computed by multiplying by a factor *k*, given by,

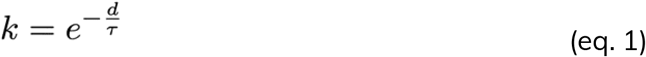

where *d* is the interval between measurements (in years) and *tau* is the decay time constant of tritium (17.75).

The dashed curve (**Fig. 14A**) was obtained in this fashion and the expected tritium decay would only account for a small fraction of the observed decline.

#### Signal-to-noise ratio

To evaluate the performance of a detector, we need to consider not only the measurements made with the light source but also the background response obtained under dark conditions. Dark currents increase with gain, vary between PMT units, and can change over the lifetime of the PMT.

Compute signal to noise ratio (SNR) at each gain setting, *g*, as the difference in mean pixel value between the light response and dark response images, divided by the standard deviation of the dark response image:

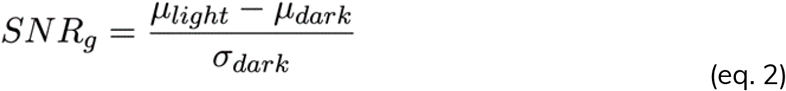

Figure 13 shows that SNR increases as a function of gain and that unit AFN9975 achieves higher SNR than the other two units. Figure 14 shows that SNR declines over time.

#### Receiver operating characteristic - area under the curve (ROC-AUC)

Although pixel grayscale values and SNR are informative, we recommend using ROC-AUC analysis as the best means to compare PMT units and track performance over time. Based on ideas from signal detection theory, ROC-AUC analysis enables comparison of the pixel value *distributions* obtained with the light source versus under dark conditions: A good detector will produce well separated distributions of the desired signal and the detector dark noise, with minimal overlap. Note that SNR partially addresses this, but does not take into account the distribution of pixel values in light response images (a product of several factors including shot noise, multiplication noise and gain).

Perform ROC-AUC analysis at each gain setting, using the distributions of pixel grayscale values in the light response image and corresponding dark response image. We provide example code for such analysis. Figure 15 shows these distributions at a variety of gain settings for an example PMT.

ROC-AUC initially increases with gain before reaching a plateau (**Fig. 15**). Moreover, although unit AFN9975 has the highest anode luminous sensitivity, highest pixel values and highest SNR, it does not have the best performance as measured by ROC-AUC. Rather, unit AFK8564 has marginally better performance. Unit AHB4783 is notably poorer, despite having similar pixel values and SNR versus the best performing unit, which is likely due to its high variance (not shown). Figure 13 shows that ROC-AUC values decrease substantially over 1.5 years of routine use.

#### Optimal gain settings

The ROC-AUC analysis can also be used to guide the choice of gain setting for imaging experiments. Specifically, the choice of gain setting should take into account:

1. ROC-AUC performance. Ideally a gain setting will be selected close to the plateau of the ROC-AUC curve.
2. HV across the PMT electrodes must not exceed the maximum value stated on the datasheet and should ideally be 20% below this value. Excessive HV can cause field emission from the dynodes and substantially shorten PMT life^62^.
3. Anode current should be kept within safe limits, typically no more than a few μA. Refer to the datasheet for specific PMT models.

Average anode current can be estimated at each gain setting using the mean pixel grayscale value of the light response image along with knowledge of the TIA and ADC settings:

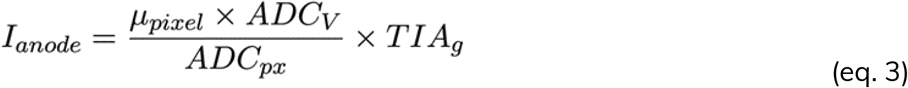

where, μ_pixel_ is the mean pixel grayscale value of the light response image at a given gain setting; ADC_V_ is the voltage that the digitiser will map to the highest grayscale value (e.g. 1 V); ADC_px_ is the corresponding grayscale value (e.g. 2048); TIA_g_ is the gain of the transimpedance amplifier (e.g. 100*10^−6^A/V).

Consider unit AHB4783 when it was first installed (**Fig. 13**, red curve). ROC-AUC increases with gain but starts to plateau at a control signal voltage of 3250 mV. The corresponding HV is safely below the limit for this PMT model and mean anode current is acceptable at around 5 uA. Thus, this gain setting would be chosen for imaging experiments.

### Estimating Absolute Magnitudes of Fluorescence Signals

#### Introduction

The overall acquired fluorescence signals are influenced by factors such as laser power, focus, properties of the fluorescent indicator, light scatter, and overall detection efficiency. It is imperative to understand and control each factor. Mismanagement of any of these factors can lead to a substantial decrease in signal intensity.

Typically, fluorescence signals are commonly expressed on relative scales, such as the dF/F ratio (the ratio to a baseline fluorescence level), or in arbitrary units, masking any degradation in signal magnitude. As a result, two laboratories following similar imaging protocols may record vastly different signal strengths for similar measurements, such as somatic calcium signals, without realising the discrepancy. Low signal magnitudes lead to noisier, less precise measurements. However, without a method to evaluate quantitative signal magnitudes that could allow a lab to detect and address imaging system problems, labs may instead attempt to only compensate for decreased magnitudes of signals through post processing, missing opportunities for improving the primary data.

We suggest reporting fluorescence signals in absolute physical units such as detected photon counts per second—instead or in addition to simply reporting the relative dF/F signal. This standardised method offers a consistent and clear way to demonstrate signal levels, making it an invaluable tool for longitudinal system performance monitoring and simplifying comparisons between different imaging systems.

Direct photon counting, while feasible, is currently rare in multiphoton microscopy, as it requires specialised electronics^65^. However, it is possible to use signal noise statistics to accurately estimate the detector *photon sensitivity* and translate detected signals into estimated photon counts. Photon count estimation is well established in photon-limited imaging modalities such as radiography^66^. Prior multiphoton studies have included variations of this procedure in their analysis ^67–69^. However, the method has not yet been applied uniformly across labs to establish quantitative benchmarks. Here we provide a step-by-step guide for estimating the photon sensitivity, the photon flux (photon counts per unit area per unit time), and photon rates (e.g. photon counts from an entire cell per unit time). We also provide a guide for interpreting these results. This approach offers a quick and intuitive way to evaluate imaging performance. However, it is important to recognise that it does not provide a detailed understanding of specific problem sources within the system. While the method works best on static fluorescent slides, it can work well even in the presence of physiological signals although care must be taken to recognize and isolate the quantum noise. The consistent application of this absolute metric can lead to universally accepted standards for recorded signal quality, fostering precise expectations and enhancing reproducibility across the scientific community.

#### Estimation of Photon Sensitivity

Light, with its stochastic, quantal nature, leads to unavoidable fluctuations in measured signal intensities, commonly termed as “shot noise,” “quantum noise,” or “Poisson noise.” An optimised microscope will function in a photon noise-limited mode, where other sources of noise have been minimised, leaving only the inevitable quantum noise as significant.

The statistical features of quantum noise follow the Poisson probability distribution, as photon detections are independent discrete events. However, additional factors like the high-frequency pulsing of laser power (typically 80 MHz in 2-photon microscopy), the PMT’s inherent gain stochasticity, and noise from the amplification process, add complexities. As a result, photons do not appear as discrete events in the signal and, for a given number of detected photons, the recorded pixel intensity will vary substantially. Additionally, the Analog-to-Digital Converter (ADC) may introduce a bias and additional non–poissonian electronic noise.

Yet, despite these complexities, the detected noise retains its essential Poissonian trait: the variance of quantum noise is linearly proportional to signal intensity whereas the slope of this linearity reveals the *photon sensitivity*, the average increase in measured light per detected photon. Other sources of noise may add on top of quantum noise but they may be recognized and isolated by their non-Poissonian traits. Finding a linear dependency between signal intensity and its variance provides a strong indication of the quantal nature of the noise. Accurately estimating the photon sensitivity allows effective translation of measured intensities into photon counts.

#### The Photon Transfer Curve

The principal tool for this estimation is the Photon Transfer Curve (PTC), which plots the variance of detected intensities against intensity values^70^. It is a remarkable fact that the relatively simple calculation of PTC provides so much insight into the properties of the image acquisition process in diverse imaging scenarios. The PTC can be derived from imaging a static fluorescent object over a short time period and estimating the noise variance across a wide range of detected intensities.

However, for convenience, we can calculate the PTC during regular experiments in the presence of physiological signals and motion. In this case, the effects of the quantum noise must be isolated from other sources of signal variance. In contrast to quantum noise, dynamic fluorescence fluctuations, such as neuronal activity or tissue movement, produce temporal variances that scale quadratically with intensity. They also have correlated spatial and temporal structures, whereas quantum noise stays decorrelated. The ability to differentiate between the poissonian quantal noise from other sources of variance enables robust estimation of photon sensitivity even during regular experiments in the presence of physiological signals and motion, without the need for a separate procedure. Estimating the PTC from the experimental data ensures consistency with acquisition settings, contributing to the efficiency of the process. For more deliberate troubleshooting, repeating the procedure with a static fluorescent preparation helps in producing more accurate results.

### Results of Photon Transfer Curve Estimation from Experimental Data

**Figure 17** presents the outcomes of PTC estimation derived directly from experimental data using a 500-frame sequence taken from the MICRONS dataset^71,72^ and following the procedure described here. The average frame is displayed in **Fig. 17A**.

**Figure 17.**
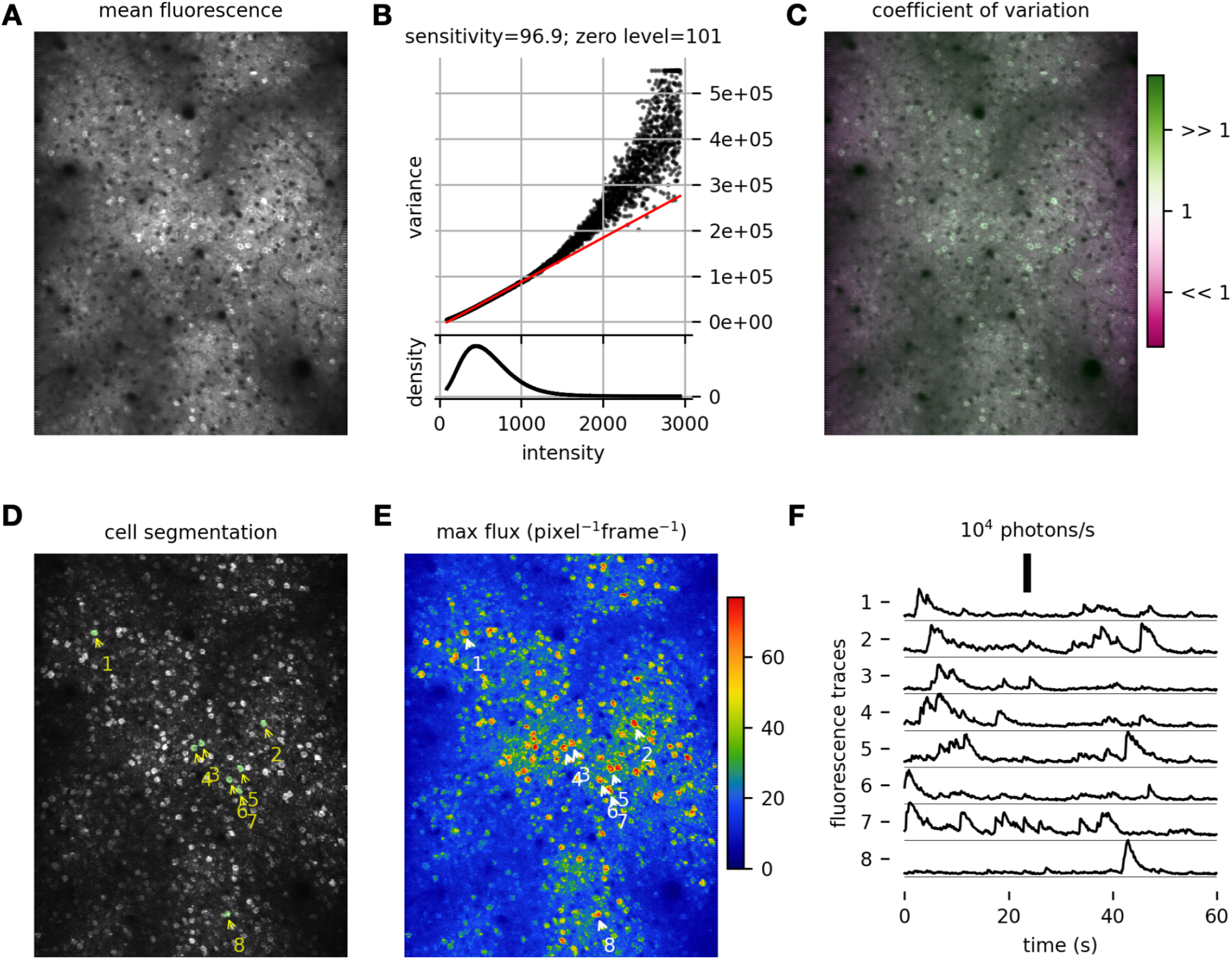
Photon transfer analysis. **A:** The average image of a 500-frame two-photon calcium imaging sequence in mouse visual cortex recorded at 8 frames per second. **B:** The Photon Transfer Curve computed from the same sequence. It features a long linear portion corresponding to poissonian noise dominating the frame-to-frame variance in all but the brightest regions. The slope of the robust linear fit (red line) reveals the photon sensitivity of 96.9 grayscale levels per photon. Note that the density of intensity values follows a long-tail distribution. The variance in bright regions of the image grows faster than predicted by the linear fit, reflecting the presence of physiological signals. Static images lack such deviations. **C:** The Coefficient of Variation image reveals areas of higher variance than predicted from quantum noise alone. Calcium activity in cells produces a higher CoV, shading them green. **D:** Cell segmentation based on the max projection image; eight cells are delineated. **E:** The maximum photon flux per pixel expressed in the units of photons per pixel per frame. **F:** The fluorescence traces from the labelled cells expressed as photons / s. The scale bar on top measures 10^4^ photons per second per cell.

#### Photon Transfer Curve

The measured photon transfer curve (**Fig. 17B**) reveals a characteristic linear portion at the lower intensity range, indicating the poissonian properties of the quantum noise. The upper intensity range is dominated by fluorescence signals such as neuronal activity and tissue motion, causing more dispersion with rapidly increasing variance. A robust linear fit (indicated by the red line) isolates the slope of the linear component, the *photon sensitivity*, with a value of 96.9, signifying that the system digitises images so that 96.9 grayscale levels are used to quantize the average intensity due to one detected photon. The intercept of the linear fit with the x-axis designates the inferred true dark level.

#### Coefficient of Variation Image

This image (**Fig. 17C**) encodes the spatial distribution of the coefficient of variation—the ratio of pixel mean values to their variances. Properly scaled by the photon sensitivity, the coefficient equals 1.0 for any Poisson process. Regions with exact Poisson noise prediction appear grey, higher variability regions appear green, and lower-than-expected variance regions appear purple. For instance, green neuronal bodies reflect added variance from neuronal activity, and purplish bands along the frame’s edges result from the acquisition system’s compensation for slower laser scanning speeds near the boundaries. Since we estimate the average photon sensitivity across the entire image, the method overestimates the photon sensitivity close to edges where the system applies a lower gain. For the same brightness, more photons are detected and less noise results, producing a lower coefficient of variation. This compensation can be undone by using a more accurate local photon sensitivity estimation, although it was not performed here where we estimate the average photon sensitivity for the entire image for simplicity.

Both the Photon Transfer Curve and the Coefficient of Variation image serve as diagnostic tools to detect diverse imaging anomalies such as pixel saturations and extra noise sources. Saturated or clipped regions, for instance, will exhibit low coefficients of variation whereas motion, laser fluctuations, and physiological signals will produce high coefficients of variation in the bright regions of the image.

#### Photon Flux Estimation

**Fig. 17D** shows several segmented cells detected by thresholding a max projection image across time, subtracting the mean fluorescence. Using the linear fit from the PTC, the entire movie can be rescaled into photon flux units, such as photons per pixel per frame, or further into physical units such as photons/μm^2^/s. **Fig. 17E** depicts the max projection image across the 500 frames expressed in units of photon flux. Note that the mean photon rates will be significantly lower. The density of intensities shown in **Fig. 17B** indicates that the majority of pixels have intensities on the order of 400, corresponding to (400 – zero level) / sensitivity = 3 photons per pixel per frame in this particular sequence.

The final result of this procedure is the conversion of fluorescence movies into the units of photon flux (photons per unit area per second). Then a separate calculation translates fluorescence traces into photon rates (photons/s) by integrating the photon flux over their image regions of interest. Measuring fluorescence in absolute units becomes valuable for monitoring signal quality across various experiments and laboratories. **Fig. 17F** shows the photon rates for the regions of interest (ROIs) corresponding to the detected cells from **Fig. 17D**. Each ROI contains 18-24 pixels with uniform weights.

### Comparison to Another Dataset

To illustrate the method, we applied the same method to a completely different dataset from another lab (**Fig. 18**). This sequence uses a different fluorescent dye, laser setting, optics, and acquisition parameters. Here, the system applies higher gains to compensate for lower pixel dwell times, producing a photon sensitivity of 678.7 grayscale levels per photon (**Fig. 18A**). While the imaging setups differ substantially, we arrive at comparable magnitudes of the fluorescence signals with peak amplitudes reaching in excess of 10^4^ photons per second from each cell (**Fig. 18D**).

**Figure 18.**
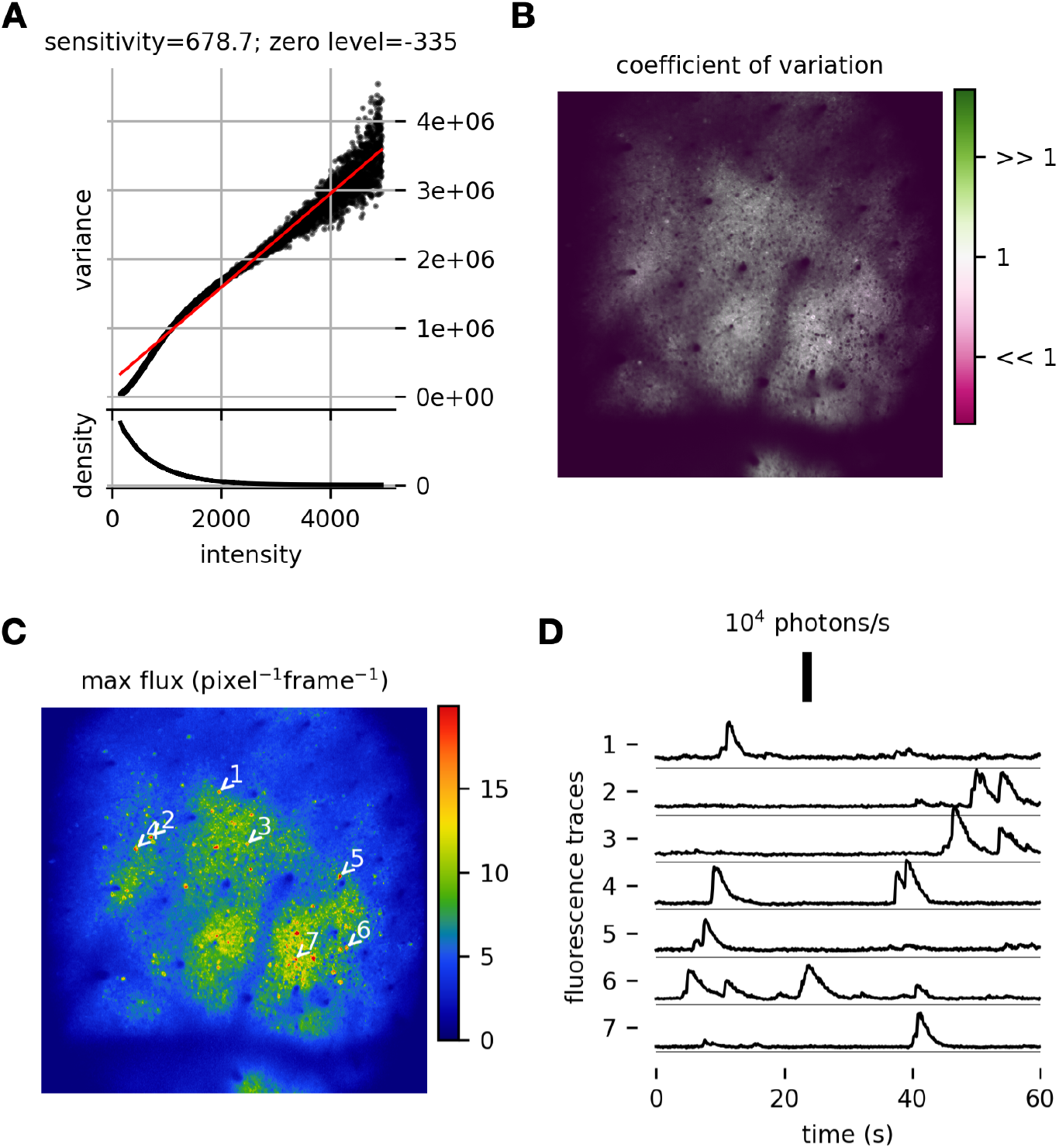
A different image sequence from another source. **A:** The Photon Transfer Curve indicates that most pixels don’t see a photon in each frame. The PTC has a non-poissonian segment where no-photon are detected. **B:** The Coefficient of Variation image reveals no deviations from predicted variance. **C:** The maximum photon flux is substantially lower than in our first dataset, due the finer pixel pitch. However, the number of pixels per cell is about 4 times larger. **D:** After ROI averaging, fluorescence signals produced comparable peak amplitudes to the first data set.

### Procedure

The following procedure estimates the photon sensitivity directly from the experimental data. For more detailed and deliberate diagnostics and troubleshooting, investigators are encouraged to use a standard static fluorescent preparation featuring a wide range of fluorescence levels, following the same procedure. The photon sensitivity is a stable property of the system since it is controlled by the PMT gain and gains applied by the acquisition system; it is insensitive to many other aspects of the imaging configuration such as magnification, the numerical aperture, laser power, etc. However, the photon transfer curve and the photon counts can describe both the quality of the fluorophore expression and the microscope’s imaging performance.

The implementation example can be accessed at the GitHub repository https://github.com/multiphoton-tools/compress-multiphoton. This implementation includes examples from a variety of projects, accessed through the DANDI Archive.

#### Calculating the Photon Transfer Curve

##### 1. Extract a Sequence

Extract a sequence (X) of approximately 500 frames from a raw imaging sequence before any processing such as motion correction or filtering.

##### 2. Equalise Photon Sensitivities (Optional, Applicable to Resonant Scanners)

Rescale the image intensity according to estimated laser dwell times at each pixel to restore uniform photon sensitivity across the image. This step reverses the gain compensation performed by the acquisition system and can aid in more accurate photon sensitivity estimations. We did not include this step in our implementation example.

##### 3. Exclude Regions (Optional)

Exclude regions near image boundaries where laser blanking and mirror vibrations might affect measurements. In our example implementation, we excluded 4-pixel margins around image boundaries.

##### 4. Calculate Mean and Difference Values

Determine the rounded mean values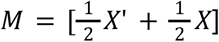 and the squared difference values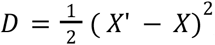 where X’ is X delayed by one frame.

##### 5. Construct Count, Intensity, and Variance Vectors

- Let vector *I* represent all unique pixel intensity values in *M*.
- Construct the count vector *C* so that each element *Cj* contains the number of pixels *k* for which *M k* = *I j*.
- Compute the variance vector *V* so that each element *Vj* contains the average value of *D k* across all pixels *k* for which *M k* = *I j*.

##### 6. Plot the Photon Transfer Curve

Plot *V* against *I* to create the Photon Transfer Curve.

Calculating means and variances from the differences between adjacent frames offers a more precise method to isolate the uncorrelated quantum noise from influences such as neuronal activity. This approach is superior to alternative methods that estimate these values over extended time periods, as it better targets the specific characteristics of quantum noise.

#### Calculating the Photon Sensitivity and Photon Flux Movies

##### 1. Estimate the photon sensitivity

Determine the photon sensitivity *q* and the zero intensity level *I*0 by performing a linear fit to the photon transfer curve so that *V* ≈ *q* · (*I* − *I*0). Ensure reliable results by weighing the fit with the pixel count vector *C* and employing a robust fitting method that minimises the impact of outliers. Our procedure utilised the Huber linear regressor from the **sklearn** package in Python^73^.

##### 2. Visualise the Fit

Plot the linear fit alongside the Photon Transfer Curve as depicted (**Fig. 16B**). Look for the characteristic linear portion where photon noise dominates. For static objects, the entire Photon Transfer Curve should align with the linear fit. In live experiments, expect a linear and tight component in darker regions and a quadratic, dispersed component in bright regions.

##### 3. Produce Photon Flux Movies

Using the fit coefficients *q* and *I*, rescale the image sequence as 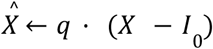 to create photon flux movies where pixel values represent photon flux in units of photon counts per pixel per frame. Alternatively, rescale the images as 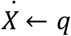 · (X − *I*_0_) /(*dx dy dt*) for units of photons per square micron per second, where *dx* and *dy*denote the pixel pitch in microns, and *dt* denotes the frame period in seconds.

#### Interpreting nonlinear Photon Transfer Curves

A Photon Transfer Curve (PTC) without a pronounced linear component *may* point to imaging issues:

##### Low Photon Rates

Extremely low photon rates can cause nonlinear PTCs. Our method can produce a nonlinear kink for rates near single photons per frame, complicating sensitivity estimation in dim images.

##### Scene Activity

Variances growing quadratically with intensity may dominate the PTC in cases of strong fluorescence signals. Repeat the procedure with a static fluorescent object if needed. However, in multiphoton imaging, we have not encountered such issues even after re-analysing diverse datasets from multiple labs.

##### Non-Poissonian Noise

Other noise sources, such as excessive electronic noise or laser fluctuations, can also result in a nonlinear PTC. These issues should be separately diagnosed and addressed to restore the imaging system to a photon noise-limited state.

### Converting Traces into Their Photon Rates

The availability of the photon flux movie 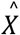, as derived above, enables the computation of fluorescence signals, such as somatic calcium traces, in terms of their absolute magnitudes, expressed as photon rates.

#### 1. Apply Motion Correction to Photon Flux Movie

Correct the photon flux movie 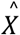 for motion. The pixel interpolations performed by the motion correction algorithm will have a negligible effect on photon rate estimation. Other processing steps, such as temporal or spatial filtrations, might have more complex effects and require careful consideration (not covered here).

#### 2. Extract Absolute Fluorescence Signal (Simple Case)

In the simplest scenario, extract the absolute fluorescence signal *y* by summing pixels over the region of *R* with equal interest weights:

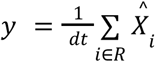

The magnitude of trace *y* will be correctly expressed in units of photons per second.

#### 3. Compute Absolute Fluorescence Signals Using a Weighted Mask

The signal *y* can also be computed using a weighted mask *h* as:

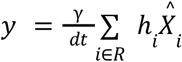

Here γ is the normalisation coefficient for proper scaling of the photon sensitivities. This computation is not trivial for dynamic scenes. The optimal unbiased scaling coefficient is:

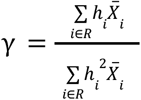, where 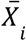is the time-averaged pixel value in 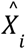

A simpler normalisation 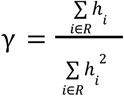 provides an accurate estimation when the image under the mask is approximately uniform. When the image is non-uniform, then this normalisation results in a lowered estimation. We recommend using this simpler normalisation.

More complex signal extraction algorithms that rely on regularised numerical methods may require a more nuanced approach to estimating the effective photon rates, which is not covered here. Our example implementation includes a notebook for deriving the scaling coefficients from first principles, which can be extended to more sophisticated methods.

## Photon Rate as a Signal Quality Metric in Multiphoton Imaging

The photon rate serves as a paramount metric for assessing signal quality from a given source. Its determination is multifactorial.

### Multiphoton Nonlinearities

Photon rate exhibits a quadratic dependency on laser power in two-photon imaging and, respectively, a cubic dependency in three-photon imaging. In two-photon imaging, the photon rate is modulated as the fourth power of the numerical aperture (NA) (Zipfel 2003). However, obstacles in the optical path or an underfilled objective lens—often resultant from suboptimal beam alignment—can substantially attenuate the effective NA.

### Laser Dwell Time

The laser dwell time on a cell will linearly amplify its photon rate. The dwell time scales with the square of the magnification.

### Imaging Depth

An exponential decline in photon rate is observed with increasing imaging depth, attributable to the scattering phenomena of excitation and collection light within tissue.

### Fluorophore Properties

The photon rate is directly proportional to both the fluorescent dye concentration and the optical cross-section of the respective fluorophore molecules.

### Optics and Detection

Variations in Group Delay Dispersion (GDD), excitation wavelength selection, optical filter specifications, and Photomultiplier Tube (PMT) detection efficiency are all contributory factors to photon rate fluctuations.

### Image segmentation

The photon rate from a cell depends on how well the cell is segmented. Including more pixels in the mask will produce higher photon rates. Beware that the photon rate calculation does not penalise for including extra pixels that are not part of the object of interest.

Importantly, the photon rate negates the influence of secondary parameters, such as PMT gain, scanning frame rate, pixels per line, lines per frame, and the grayscale parameters of the ADC circuitry and the acquisition software. It is important to ensure that no spatial or temporal filtration is applied to the images before the photon rate estimation since it can bias the estimation.

### Signal Quality Awareness

In the course of employing this technique, we have observed significant disparities (orders of magnitude) in photon rates across datasets sourced from various laboratories, even under ostensibly uniform imaging parameters. Such divergences, more often than not, go unnoticed by the scientific community, making it difficult to pinpoint the underlying causes until further collaborative scrutiny. Defining concrete benchmarks for photon rates under particular conditions may be premature. Nevertheless, the uniform quantifications of absolute signals will engender progressive shifts in both experimental paradigms and scientific communication.

### Strategies for Enhancing Photon Rates

On occasions, low photon rates, especially in somatic calcium signals, might be expected, particularly in deeper tissue imaging or large fields of view. However, improvements can be achieved by multiple approaches: the incorporation of brighter fluorescent indicators, increased laser power, optimization of excitation wavelengths, replacement of aged PMTs, or improved beam alignment and the GDD (Group Delay Dispersion) parameters.

## Author contributions

Investigation: **All authors**

Writing – original draft:

Laser Power at the Sample: **R.M.L**., **R.A.A.C**., **N.O**.

Field of View Size: **R.A.A.C**., **N.O**.

Field of View Homogeneity: **R.A.A.C**., **N.O**., **R.M.L**.

Spatial Resolution: **S.L.S**., **C-H.Y**.

Pulse Width Control and Optimisation: **D.S.P, R.A.A.C**.

Photomultiplier Tube Performance: **I.H.B**., **B.P**.

Estimating Absolute Magnitudes of Fluorescence Signals: **D.Y**.

Writing – review & editing: **All authors**

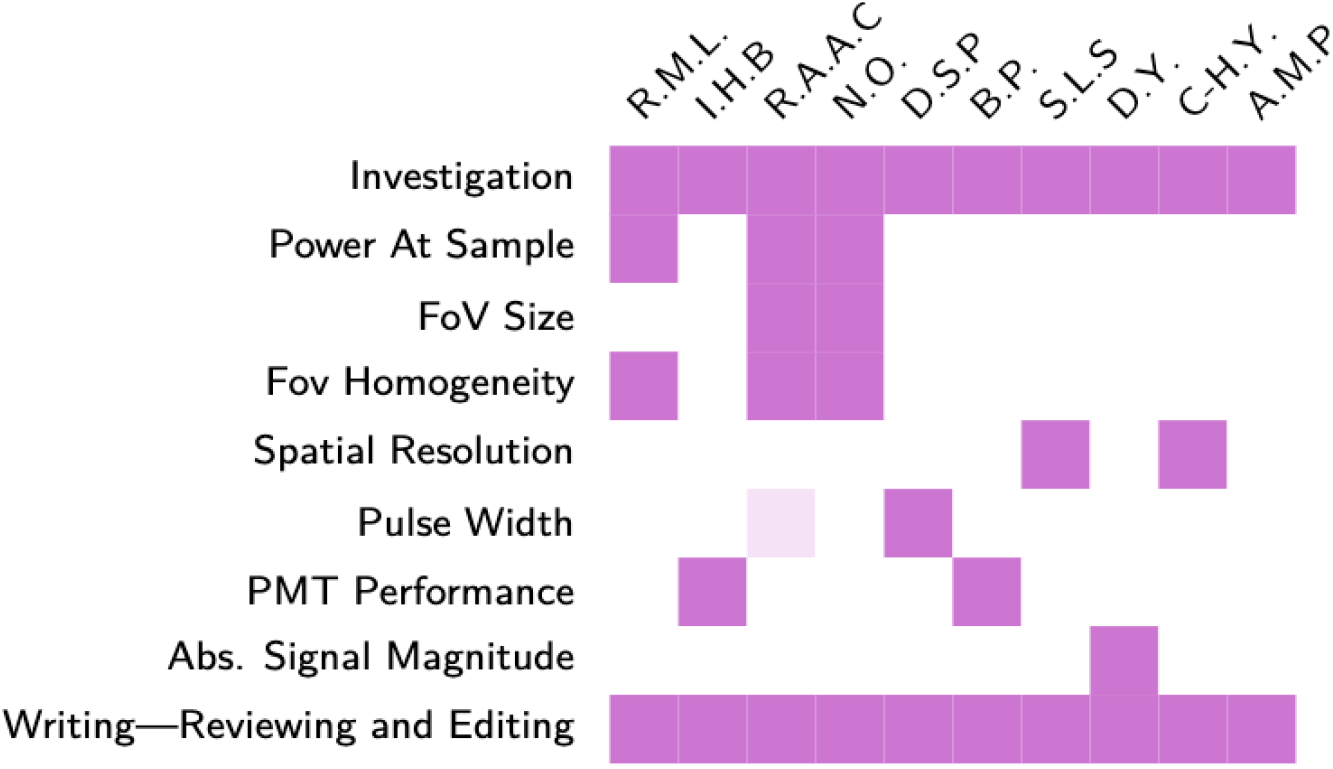

## Acknowledgements

This work was supported by funding from the Wellcome Trust (204651/Z/16/Z, 220273/Z/20/Z), the European Research Council (ERC) under the European Union’s Horizon 2020 research and innovation program (grant agreement No 852765), the National Institutes of Health (R01EY035378 and RF1NS121919 to S.L.S., U24NS116470 to D.Y., U19NS104649 and U01NS113273 to D.S.P.), and the National Science Foundation (1934288 to S.L.S.).

## Conflicts of interest

R.M.L., I.H.B., R.A.A.C., N.O., D.S.P., B.P., S.L.S., D.Y., C-H.Y., A.M.P. have no conflicts to declare.

## Notes

### Competing Interest Statement

The authors have declared no competing interest.

